# Genomic Consequences of Altered Flood Regimes: Gene Flow and Adaptation in a Threatened Floodplain Plant

**DOI:** 10.64898/2026.09.28.755008

**Authors:** Wen-Hsi Kuo, Eduardo Aguirre-Mazzi, Burgund Bassüner, Matthew A. Albrecht, Ethan Coble, Brad Delfeld, Noah Dell, Megan J. Engelhardt, Jordan Hathaway, Maia M. Jones, Rebecca Sucher, Sage Waite, Maia Washington, Christine E. Edwards

**Affiliations:** Missouri Botanical Garden; Washington University in St. Louis; Appalachian State University; University of West Alabama; Dickinson College

**Keywords:** Gene flow, Fitness, Genetic load, Adaptive trait, genomics, Floodplain

## Abstract

Floodplains are highly dynamic ecotones where species persistence depends on synchrony between natural flow regimes and life-history. Artificial river engineering has profoundly altered these systems by reducing the frequency and magnitude of spring floods while increasing the unpredictability of flood timing and intensity during summer. These hydrological changes have disrupted historical patterns of dispersal, growth, reproduction, and competitive success among floodplain specialists. Effective conservation in these ecosystems therefore requires understanding how human-altered flow regimes have reshaped floodplain specialists’ demography, genetic diversity, genetic connectivity, functional variation, and adaptive potential. Here, we address this challenge in *Boltonia decurrens*, a federally threatened floodplain plant endemic to the Illinois River Valley, by integrating a chromosome-scale reference genome, whole-genome resequencing, and common-garden adaptive trait measurements from 424 individuals across 17 populations. Demographic analyses revealed a severe recent bottleneck and a subsequent rebound. Population structure was clinal with downstream-dominant gene flow, indicating that regional population connectivity predominates over longer-distance gene flow. A subset of populations showed elevated genetic isolation with increased inbreeding and homozygous deleterious burden, revealing a genomic cost of disrupted floodplain connectivity. Common-garden observations and GWAS showed extensive within-population phenotypic and genetic variation in flowering time, flowering stem height, and year of first flowering (1^st^ vs. 2^nd^ year), which are hypothesized to buffer populations and to be selectively maintained under human-altered flow regimes. Finally, landscape-genomic approaches detected a divergent haplotype block separating northern and southern populations, suggesting that adaptive divergence can persist even in a system shaped by recurrent disturbance and ongoing gene flow.

## Introduction

Floodplain and riparian habitats are dynamic ecotones between aquatic and terrestrial ecosystems, characterized by strong spatial and temporal heterogeneity in hydrology, sediment dynamics and disturbance (Naiman & Décamps, 1997; Tockner & Stanford, 2002; Ward, Tockner, & Schiemer, 1999; Wohl, 2025). This heterogeneity generates shifting mosaics of microhabitats supporting plant communities with diverse functional strategies, making riverine floodplains among the most species-rich environments globally (C. Blom, 1999; Tockner & Stanford, 2002; Ward et al., 1999). Floodplain environments are maintained by seasonal flooding, which helps to maintain open, high-light habitats, limiting competitive dominance by later-successional or generalist species, with nutrient pulses for recruitment (Dewoody, Nason, & Smith, 2004). For floodplain specialists, persistence depends on dispersal-mediated connectivity and recolonization following periodic extirpation, as well as adaptive trait variation that enables populations to withstand and recover from recurrent variation in flood timing, frequency, depth and duration (C. W. P. M. Blom et al., 1990; C. W. P. M. Blom & Voesenek, 1996; C. W. P. M. Blom et al., 1994; Lytle & Poff, 2004).

Human river engineering activities, including channelization, levee and dam construction, flood suppression, and land conversion, have disrupted the hydrological connectivity and natural flow regimes that structure many floodplains, leaving formerly connected floodplain systems fragmented and degraded (Tockner & Stanford, 2002; Ward et al., 1999; Wohl, 2025). Reduced connectivity can increase inbreeding and genetic erosion and prevent recolonization after local extirpation. Altered natural flow regimes may decouple the synchronization of life history events with environmental conditions and increase competitive pressure from later-successional or habitat-generalist species (Dewoody et al., 2004; Hevroy, Moody, & Krauss, 2018; Lytle & Poff, 2004; Murray, Reid, Capon, & Wu, 2019). Climate change further compounds these pressures by increasing the frequency of irregular and extreme precipitation, intensifying drought and flooding, and altering nutrient pulses (Bian et al., 2026; Smith, Caswell, & Mettler-Cherry, 2005). Finally, the altered environmental conditions caused by anthropogenic activities can bring closely related species into greater contact, increasing the potential for interspecific gene flow, which can sometimes negatively affect rare species (Grabenstein & Taylor, 2018; Rhymer & Simberloff, 1996). However, few studies of floodplain plants have assessed how both past and present environmental conditions have shaped inter- and intraspecific variation and gene flow, genetic erosion, and adaptive trait variation. Such studies can help guide conservation strategies for sustaining biodiversity and ecosystem function in engineered riverine landscapes (Alberto et al., 2013; Dauphin et al., 2023; Hoelzel, Bruford, & Fleischer, 2019; Teixeira & Huber, 2021).

Here, we use *Boltonia decurrens* (Torr. & A. Gray) A. Wood (Asteraceae: Boltoniinae) as a model floodplain herb, integrating phylogenomics, population genomics, and functional trait analyses to inform conservation. *Boltonia decurrens* is endemic to a ~400 km reach of the Illinois River Valley and its confluence with the Mississippi River (Dewoody et al., 2004; Nesom, 2000; Smith, Keevin, Mettler-Mcclure, & Barkau, 1998). As an early-successional floodplain specialist, *B. decurrens* depends on annual spring flooding both for hydrochorous seed dispersal and for the creation of open, sandy alluvial substrates with nutrient pulses required for germination and seedling establishment (Smith, Brandt, & Stone, 1995; Smith et al., 1998; Smith & Keevin, 1998; Smith & Moss, 1998). Over the last century, river engineering, including construction of levees for flood control and locks and dams for navigation, has reduced the annual spring flood, disconnected floodplains, and diminished both habitat quality and availability; this contributed to the near disappearance of *B. decurrens* from the wild before its listing under the U.S. Endangered Species Act in 1988 (USFWS; 53 FR 45858). However, it is unclear how changes in the past and present flood regimes, population size, habitat quality, and connectivity have shaped genetic diversity and structure in *B. decurrens*.

Although *B. decurrens* is narrowly distributed in floodplain habitats, its congener *B. asteroides* complex (hereafter, *B. asteroides*) is broadly distributed across eastern and central North America, occupying a wider range of mesic to hydric habitats. Although the two species are partially sympatric, previous studies found hybridization and introgression between them was uncommon; even though no strong physical barriers to gene flow separated these taxa (DeWoody, Nason, & Smith, 2011). However, anthropogenic activities such as habitat restoration may bring these species into increased contact, potentially increasing the likelihood of interspecific hybridization (Grabenstein & Taylor, 2018; Rhymer & Simberloff, 1996). Furthermore, because previous analyses employed allozyme markers and the detection of interspecific gene flow may be limited by the resolution of the genetic markers, additional analyses with genomic-scale markers are needed to assess the relationship between *B. decurrens* and other *Boltonia* species, the extent of interspecific hybridization and introgression, and the possible consequences for the conservation of *B. decurrens*.

New challenges, such as increasing variability in the timing of floods, may impose novel selection on life-history strategies in *B. decurrens* that affect their response to catastrophic stresses (Bian et al., 2026). For example, *B. decurrens* was previously found to exhibit variable life histories ranging from annual to biennial and short-lived perennial forms. Previous authors hypothesized that annual spring flooding would favor an annual life history and delayed or sporadic floods would favor biennial or perennial strategies (Smith et al., 2005). This combination of flood-dependent recruitment, variable life history, metapopulation dynamics, and recent habitat alteration make *B. decurrens* well-suited for evaluating the processes shaping adaptation in human-altered floodplain landscapes.

Documenting local adaptation has become a priority in conservation biology because it can help delineate conservation units, develop localized management plans, and inform future assisted migration if needed (Aitken & Whitlock, 2013; Funk, McKay, Hohenlohe, & Allendorf, 2012). Although the dynamics of local extirpation and recolonization of the frequently disturbed environments occupied by *B. decurrens* would not necessarily favor the development of strong local adaptation, the 400-km Illinois River Valley spans a marked temperature range, which could favor divergence in life-history strategies, such as flowering phenology. More broadly, hydrology, nutrient availability, and sediment composition may also shift along the river corridor through the input of tributaries and the influence of major confluences. In particular, populations in the Illinois River Valley occur on glacial outwash and sandy alluvial sediments, whereas those that occur near the confluence with the Mississippi River occupy more clay-rich soils (Smith et al., 1995). However, it is unclear whether *B. decurrens* shows local adaptation to this environmental variation.

In this study, we used *B. decurrens* to ask how connectivity, fragmentation, and selection have interacted to influence interspecific hybridization, genetic diversity, genetic structure, and adaptation of a threatened floodplain plant. We first evaluated connectivity at two evolutionary scales: among species, by testing whether *B. decurrens* represents a distinct conservation lineage and whether hybridization or introgression with sympatric congeners such as *B. asteroides* contributes to contemporary genomic variation at the population level. We then examined how population connectivity or isolation have influenced genetic diversity, inbreeding and genetic load, and how heterogeneous floodplain environments have shaped adaptive traits associated with establishment and reproduction. Finally, we asked whether local adaptation could have emerged and persisted in a system characterized by high gene flow, recurrent disturbance, and metapopulation turnover. Together, these questions allow us to evaluate how interspecific and intraspecific population connectivity, adaptive trait variation, and genetic erosion jointly shape the persistence of a threatened floodplain species in an engineered riverine landscape.

To address these questions, we integrated *de novo* genome assembly and whole-genome resequencing, common-garden phenotyping, phylogenomics, population and landscape genomics, and genome-wide association analysis (GWAS) in *B. decurrens* and its congeners to develop an evolutionarily informed basis for conservation. <u>First</u>, we reconstructed the phylogeny of the genus *Boltonia* with complete species sampling to clarify species boundaries and assess whether historical or ongoing introgression affects its status as a distinct conservation unit. <u>Second</u>, we characterized species-wide demographic history, patterns of genetic structure and among-population gene flow to evaluate its current evolutionary status and test whether hydrochorous dispersal and riverine connectivity shape genetic structure along the Illinois River. <u>Third</u>, we characterized genome-wide patterns of diversity, inbreeding, and genetic load within *B. decurrens* populations to evaluate whether populations have experienced declines due to fragmentation and reduced connectivity. <u>Fourth</u>, we combined common-garden phenotyping with GWAS to ask whether life-history strategies, such as flowering time and stem architecture, are under heterogeneous selection in floodplain environments. <u>Finally,</u> we integrated landscape genomic approaches to identify candidate loci and structural variants of local adaptation and assess how well genomic signals of local adaptation align with the loci associated with adaptive trait variation. Together, these aims provide a functionally informed, genome-enabled framework for conserving threatened floodplain plant species in human-altered, dynamic landscapes.

## Materials and Methods

### Plant materials, plant propagation, and phenotyping

Between 1997-2011, a comprehensive survey of *B. decurrens* was conducted by Marian Smith, who collected seeds from 17 natural populations spanning the Illinois River (Dewoody et al., 2004). Within each population, multiple spatially separated maternal plants were sampled, and deposited in the Missouri Botanical Garden Seed Bank, where they were dried to 20 ± 5% relative humidity and stored at −20 ± 3°C. Metadata are available in the Missouri Botanical Garden’s living collection database (https://livingcollections.org/home).

In January 2024, we established a greenhouse common garden to quantify phenotypic variation among populations. Depending on the number of seeds available, we selected between 6-52 seeds from each of 300 maternal lines, representing all 17 collection sites. We stratified the seeds in cold/wet conditions in a dark growth chamber at 4°C for 6 weeks. In February, seeds were transferred to agar medium and placed in a growth chamber at 30/15°C with 12-hr light/dark cycles for germination. In March, with the goal of growing 1–4 individuals per maternal line, we transferred up to four germinated seeds (with visible radicles) per maternal line into seed propagation trays. Not all maternal lines produced viable seedlings. The final common garden comprised 458 individual plants from 242 maternal lines (**Supplementary File 1**). Maternal lines were randomized across trays to minimize positional effects. In mid-March, when most seedlings had primary leaves, they were transferred to a greenhouse at Oertli Hardy Plant Nursery. After approximately one week of acclimation under the mist bench, they were transferred to 127×127-mm square pots containing BM5 substrate (Berger, Saint-Modeste, QC, Canada). Temperature and watering regimes were managed to minimize environmental heterogeneity among individuals.

For phenotyping, in 2024, we measured vegetative traits (number of stems, leaf width, and leaf length) at weekly intervals and reproductive traits (number and developmental stages of capitula, flowering stem length) at two-day intervals. In 2025, we recorded the day of the first mature capitula, defined as anthesis of any disk flowers in a capitulum, the number of flowering stems produced per individual, and flowering stem length at the same date. When the apical meristem developed into a terminal capitulum, the flowering stem length was measured, after which no further growth was detected.

### Sampling, DNA sequencing, and assembly

We sampled leaf tissue for DNA sequencing. Because 10 seedlings did not survive, we sampled leaf tissue from 448 of the 458 original plants. For congeneric species and outgroups, we sampled leaf tissue from 25 specimens from the Missouri Botanical Garden Herbarium. Information for all DNA samples is provided in **Supplementary File 1**. See **Supplementary** I**nformation** for details on DNA extraction for whole-genome assembly and resequencing.

First, we *de novo* assembled a haplotype-phased and chromosome-scale reference genome of one sample of *B. decurrens* based on PacBio HiFi reads (88×) and a HiC contact map (See **Supplementary** I**nformation**). Then, population-scale samples were shotgun sequenced using 2×150 bp reads on an Illumina NovaSeq X plus to >10× coverage depth. In total, we successfully resequenced 424 individuals, representing 1-3 individuals from each of 242 maternal lines. Genotypes were mapped against the reference genome and called using the GATK pipeline. We generated multiple nuclear variant-site datasets with different combinations of samples and filtering criteria, depending on the analysis type (See **Supplementary** I**nformation**). The plastome was also assembled by GetOrganelle (Jin et al., 2020), and the same GATK pipeline was used for the population-level plastome data.

### Phylogeny reconstruction, hybridization, and introgression

To assessed species boundaries and interspecific hybridization, we reconstructed the phylogeny and plastid haplotype network. We used concatenated nuclear SNPs for maximum likelihood phylogeny inference, employing the GTR+ASC+G model to correct for ascertainment bias. Node support was evaluated with 1,000 ultrafast bootstrap replicates and 1,000 SH-aLRT replicates with `bnni` enabled to minimize bootstrap support inflation in IQ-TREE v3.0.1 (Nguyen, Schmidt, von Haeseler, & Minh, 2015). A plastid haplotype network was reconstructed and visualized in HapNetworkView using a modified TCS algorithm (Chi et al., 2025).

Using the nuclear dataset, admixture graph inference and ABBA-BABA tests were conducted to assess historical interspecific hybridization using the R package `Admixtools` (Maier et al., 2023). We also assessed admixture with a nuclear dataset including *B. decurrens* and its close relatives, *B. asteroides* and *B. diffusa*, using ADMIXTURE v1.3.0 with K = 1-20 (Alexander, Novembre, & Lange, 2009). To identify loci that show signatures of positive selection in *B. decurrens* relative to *B. asteroides*, we conducted divergence scans comparing species-wide estimates of F_ST_, nucleotide diversity (π) and Tajima’s D for *B. decurrens* and *B. asteroides* in 10-kb non-overlapping genomic windows using VCFtools. Only windows containing more than 100 SNPs were retained. Genomic regions under significant positive selection in *B. decurrens* were defined as those showing the top 0.1% of the highest F_ST_, the highest ratio of nucleotide diversity (*B. asteroides*/*B. decurrens*), and the lowest Tajima’s D values. These metrics were also compared between *B. decurrens* and *B. asteroides* to understand how rarity has affected genetic diversity.

### Demographic history

Recent species-wide effective population size (N_E_) trajectories were inferred for *B. decurrens* using GONE2 (Santiago, Köpke, & Caballero, 2025). To address non-independence among individuals caused by sampling siblings (i.e., seeds from the same maternal plant), we remove one sample from each pair with a kinship coefficient exceeding 0.2 using PLINK2 (--king-cutoff 0.2) (Chang et al., 2015). After filtering, 235 samples and 1,863,350 biallelic sites remained for demographic inference. We ran GONE2 assuming unphased diploid genotypes (-g 0), a constant recombination rate of 1.1 cM/Mb (-r 1.1), and treating the sample as a random draw from a structured metapopulation (-x).

### Intraspecific population structure and gene flow

To assess genetic structure in *B. decurrens*, we assessed the ancestral genetic composition of each individual using ADMIXTURE v1.3.0 with K = 1-20 using a nuclear data set that included only *B. decurrens* (Alexander et al., 2009). Principal component analysis (PCA) and pairwise F_ST_ estimation were conducted by PLINK2 (Chang et al., 2015). Isolation-by-distance (IBD) and isolation-by-environment (IBE) were evaluated using Mantel tests implemented in the R package `vegan` (Dixon, 2003). For IBE, climatic variables were standardized to a mean of zero and a standard deviation of one, and pairwise environmental distances among populations were calculated as Euclidean distances.

We evaluated three alternative migration hypotheses in fastsimcoal2 v2.8 (Excoffier et al., 2021) to reflect hydrochorous dispersal along the river network as a directed stepping-stone process—bidirectional, upstream-only, and downstream-only. The three models were identical except for directionality constraints in the initial migration matrix. Gene flow was permitted only among one-step (adjacent) and two-step (next-nearest) connections (e.g., one-step A↔B and two-step A↔C), with all other migration entries fixed to zero. The bidirectional model estimated directional rates for each permitted link, whereas the upstream-only and downstream-only models retained only the lower-triangular or upper-triangular entries, respectively, of the same sparse stepping-stone migration matrix (setting the opposite triangle to zero). Panmictic migration was allowed only before the divergence time TDIV. All models used six demes (A–F; see Results) with haploid effective sizes (N_A–N_F), initial values of 106, 70, 234, 90, 144, and 164 haploid genomes for A–F (actual sample size times two), and all growth rates = 0, with divergence specified by five split events at TDIV. Priors were defined in a fastsimcoal2 parameters/rules file: N_A–N_F were assigned uniform priors of 100–100,000 and TDIV a uniform prior of 10–200 generations, assuming that the annual spring floods before 1930 could cause panmictic gene flow (Mettler, Smith, & Victory, 2001; Smith et al., 2005). To reduce the number of free migration parameters while retaining the stepping-stone structure, we used two log-uniform hyper-parameters for the number of migrants per generation (NMIG_1POP: 1×10^−2^ to 20; NMIG_2POP: 1×10^−3^ to 2) and converted them to directional per-generation migration rates via mi_→j_ = N_migrants_/N_i_, generating one-step and two-step rates used in the migration matrices. Parameter estimation was conducted using maximum composite-likelihood optimization (-m -0 -C 10 -n 400000 -L 100 -s 0 -M), using minor-allele SFS likelihood, excluding the monomorphic class, ignoring SFS entries with counts <10, and estimating expected spectra from 400,000 simulations per ECM cycle across 100 ECM cycles. For each model, we conducted 100 independent maximum-likelihood optimizations to reduce the risk of local optima and to identify the global best-fitting parameter set. The parameter combination achieving the highest composite likelihood was retained as the best estimate. We then fixed parameters to this best-fit model and re-ran fastsimcoal2 100 additional times (-m -0 -C 10 -n 200000 -s 0) to generate a replicate-based likelihood distribution for the inferred model fit.

### Inbreeding, genetic diversity, genetic load

To assess inbreeding, runs of homozygosity (RoH) were identified for each individual using `bcftools roh` with -G 30 (Danecek et al., 2021). Only RoH segments ≥40 kb were retained for downstream analyses. Population nucleotide diversity (π) and Tajima’s D were calculated for each population in 10-kb non-overlapping windows using VCFtools (Danecek et al., 2011). Population-level values were summarized as the median across windows. For Tajima’s D, only windows containing more than 100 SNPs were retained. To estimate genetic load, we used SIFT (Sim et al., 2012), comparing protein-coding variant sites to orthologous sequences in NCBI RefSeq and classifying a mutation as deleterious when the corresponding amino acid state was not observed among orthologs in the database. Total genetic load (counts of total deleterious alleles per individual), heterozygotic genetic load, and homozygotic genetic load were calculated for each population of *B. decurrens* following Fiscus, Aguirre-Liguori, Gaut, and Gaut (2025).

### The genomic basis of adaptive traits and climate adaptation

We estimated narrow-sense heritability (*h*^*2*^) for each continuous trait using a genomic “animal model” with a realized genomic relationship matrix (GRM) generated by PLINK2 (Chang et al., 2015). For each trait, we fit a linear mixed model in the R package `sommer`: *Y* = *Xβ + a + e*, where *a* is the additive genetic effect and *e* is the residual error, with V*a*r(*a*) = *σ*^*2*^_*A*_*G* and Var(*e*) = *σ*^*2*^_*E*_*I*, where *G* is the genomic relationship matrix and *I* is the identity matrix. Narrow-sense heritability was calculated *as h*^*2*^ = *σ*^*2*^_*A*_ */* (*σ*^*2*^_*A*_ *+ σ*^*2*^_*E*_). To identify genetic loci associated with variation in adaptive traits, we performed genome-wide association analysis (GWAS) using PLINK2’s glm function (Chang et al., 2015). The regression models included the first four genetic principal components (PC1–PC4) as covariates to control for population structure.

To assess local adaptation across the environmental gradient occupied by *B. decurrens*, we conducted genome-wide population differentiation and environmental association analyses. We first conducted analyses in BayPass v2.31 (Gautier, 2015) using allele-count input files for 17 populations. To account for shared demographic history, we first estimated the population covariance matrix (Ω) from a reduced, high-LD–filtered dataset (329,108 SNPs) by running BayPass in Basic (no-covariate) mode. We then used this pre-estimated Ω (-omegafile) in downstream genome-wide scans based on 6,950,268 SNPs. Population differentiation was quantified using SNP-wise XtX in Basic mode. Genotype–environment associations were tested in covariate mode with two population-level covariates, which estimated regression coefficients (β) and measured support for association while controlling for Ω. MCMC settings were: 5,000 burn-in iterations, followed by 1,000 posterior samples saved with a thinning interval of 20, and up to 20 pilot runs of 500 iterations each to tune proposal parameters toward targeted acceptance rates (0.25–0.40).

To corroborate BayPass signals with approaches that do not require a priori population definitions, we performed complementary genome scans in R using LFMM2 (latent factor mixed models; LEA package) (Caye, Jumentier, Lepeule, & Francois, 2019) and pcadapt (PCA-based outlier detection) (Privé, Luu, Vilhjálmsson, & Blum, 2020). LFMM2 was used to test SNP– environment associations while controlling for cryptic structure via K = 5 latent factors, and pcadapt was used to detect loci excessively associated with population structure using K = 5 retained principal components; in both cases, K was selected from the scree plot to capture the major axes of genetic variation.

Candidate genes underlying adaptive traits were identified by focusing on loci identified by multiple genome-scan approaches (i.e., intersecting outlier signals across methods). For each intersecting signal, we extracted annotated gene models located within a ±10 kb flanking window around the focal variant sites. Putative functions were assigned to the sites using sequence similarity searches against the UniProt database (BLAST). For each gene, we recorded the best-supported UniProt matches and their associated functional descriptions.

## Results

### *De novo* genome assembly

We generated a highly contiguous, accurately phased, and chromosome-scale *Boltonia decurrens* genome assembly (**Supplementary Table S1, Fig. S1A**). The Hi-C contact heatmap showed accurate long-range scaffolding and well-resolved chromosome structure (**Supplementary Fig. S1B**). The k-mer modeling using PacBio HiFi reads (k = 21) estimated a haploid genome size of ~431 Mb and moderate heterozygosity (~1.6% homologous genome copies differ at a nucleotide position); the estimated coverages of homozygous and heterozygous regions were 88× and 43×, respectively (**Supplementary Fig. S1C**). Whole-genome alignment between haplotypes revealed extensive synteny and collinearity while retaining expected allelic and structural differences (**Supplementary Fig. S1D**). Assembly accuracy and completeness were supported by k-mer–based evaluation using an independent Illumina dataset (Merqury QV = 38.62–38.75; combined QV = 38.68; k-mer completeness = 81.55– 81.84% per haplotype and 99.32% combined) (**Supplementary Fig. S1E; Table S1**) and by high gene-space completeness (BUSCO eudicots_odb10: 95.6% and 95.4% complete; embryophyta_odb10: 97.7% and 97.5% complete for primary and alternative haplotypes, respectively) (**Supplementary Table S1**). The assemblies were predominantly assigned to chromosomes (97.69–98.88%), with identical GC content (34.46%) and broadly consistent repeat composition (57.54% vs. 57.09%; including abundant retroelements and LTRs) (**Supplementary Table S1**).

### Phylogeny, hybridization, introgression

Using complete sampling of all six described species, the nuclear phylogeny reconstructed *B. decurrens* as a well-supported, monophyletic lineage (**Fig. 1A**). The *B. decurrens* clade was sister to a paraphyletic group including several samples labeled as *B. diffusa* or *B. asteroides* Then, *B. montana, B. apalachicolensis*, and *B. caroliniana*, were placed as successive sisters to the *B. decurrens*-*B. diffusa*-*B. asteroides* clade, respectively. For all subsequent analyses, we used *B. asteroides* as the sister species to *B. decurrens* based on its dominance and wide distribution. These species relationships were similar to those found in both PCA (**Fig. 1E**) and the plastome tree (**Supplementary Fig. S2**).

**Figure 1.**
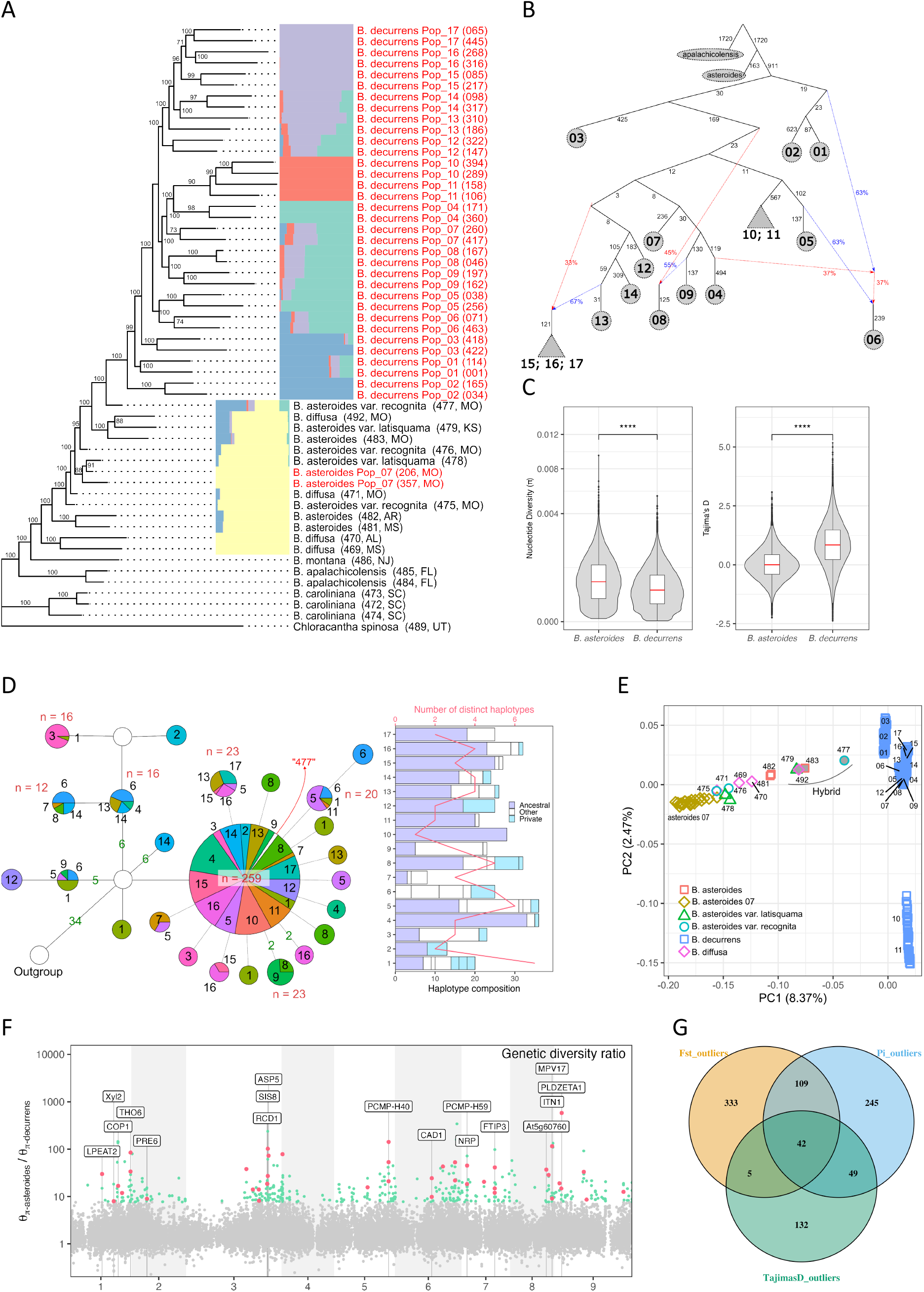
Evolution of *Boltonia decurrens* and congener species. **(A)** Maximum-likelihood phylogenetic tree of *Boltonia* spp. Bootstrap (> 70%) of branch support was labeled. The tip labels in red were sampled within the Illinois River Valley at one of the 17 *B. decurrens* populations. For visualization purposes, only two samples from each population are shown. Ancestry estimates from ADMIXTURE at K = 5 are shown to the right of the tree. **(B)** Admixture graph showing the inferred driV distance among *B. decurrens* populations and from *B. asteroides* to *B. decurrens*. **(C)** Comparison of genome-wide genetic diversity and Tajima’s D in *B. asteroides* and *B. decurrens*. **(D)** Plastome haplotype network. Unique combinations of colors and numbers represent population indices, and numbers on branches indicate mutational steps between haplotypes if steps > 1. Number of individuals of a specific haplotype is labeled if n > 10. “477”, the hybrid individual – *B. asteroides* var. *recognita* (477, MO). See **Supplementary Fig. S2** for the full plastome network including all species. **(E)** PCA of *B. decurrens, B. asteroides, B. diffusa*, and putative hybrid/introgressed individuals. Gray filled symbols denote putative hybrids. *B. decurrens* individuals are labeled by a two-digit population index, whereas individuals of other species are labeled by three-digit genotype accessions. **(F)** Genome scan for positive selection (10 kb windows) in *B. decurrens* relative to *B. asteroides*. Green points show the windows in the top 0.1% of the distribution. Red points highlight windows in the top 0.1% intersection of the genetic diversity ratio, F_ST_, and Tajima’s D scans. See **Supplementary Table S2** for the description of the candidate genes. **(G)** Venn diagram showing the overlap in outlier variant sites identified by scans of genetic diversity ratio, F_ST_, and Tajima’s D. **See Supplementary Fig. S4** for full genome-wide plots of each statistic.

Admixture analyses revealed a few putative hybrid individuals; all identified as *B. asteroides* or *B. diffusa* and collected downstream of *B. decurrens’* current distribution. These hybrids shared various degrees of ancestry with the southern populations of *B. decurrens* (population #1, #2, #3) (**Fig. 1A, E**). The southern *B. decurrens* populations also showed less distance in drift units from *B. asteroides* (**Fig. 1B**). We tested whether the partially shared ancestral composition between the southern population and *B. asteroides* was a result of incomplete lineage sorting (ILS) or gene flow (i.e., introgression); the ABBA–BABA test detected no excess derived allele sharing between *B. asteroides* and the southern *B. decurrens* populations relative to other non-southern populations (**Supplementary Fig. S3A**), but it detected excess derived allele sharing between the putative hybrids and *B. decurrens* relative to *B. asteroides* (**Supplementary Fig. S3B**).

Divergence scans identified a small subset of loci with putative signatures of positive selection in *B. decurrens* relative to *B. asteroids* across analyses of genetic diversity ratios, F_ST_, and Tajima’s D (**Fig. 1F, G**). The candidate genes were associated with cell structure, light signaling, stress tolerance, photosynthesis, and developmental regulation (**Supplementary Table S2**). At the species scale, *B. decurrens* exhibited reduced genome-wide diversity and generally positive Tajima’s D relative to *B. asteroides* (**Fig. 1C**), consistent with a reduced effective population size. Plastome haplotype relationships further revealed a broadly shared ancestral haplotype across nearly all *B. decurrens* populations (**Fig. 1D**), consistent with a demographic history involving a severe bottleneck followed by recent expansion.

### Demographic history

Analyses to quantify the timing and magnitude of recent demographic changes in *B. decurrens* over ~150 generations showed a pronounced bottleneck in *B. decurrens* approximately 12–13 generations ago (**Fig. 2G**). Assuming an average generation time of 1.5 years, this bottleneck corresponds to ~1975, predating the listing of *B. decurrens* as threatened in 1988. The inferred rebound in effective population size is also consistent with historical records of population expansion following major Illinois River floods in 1993 and 1995 (Smith et al., 1998). Together with the dominance of a single plastome haplotype across populations (**Fig. 1D**), these results suggest that much of the population structure and demographic signal documented here formed relatively recently.

**Figure 2.**
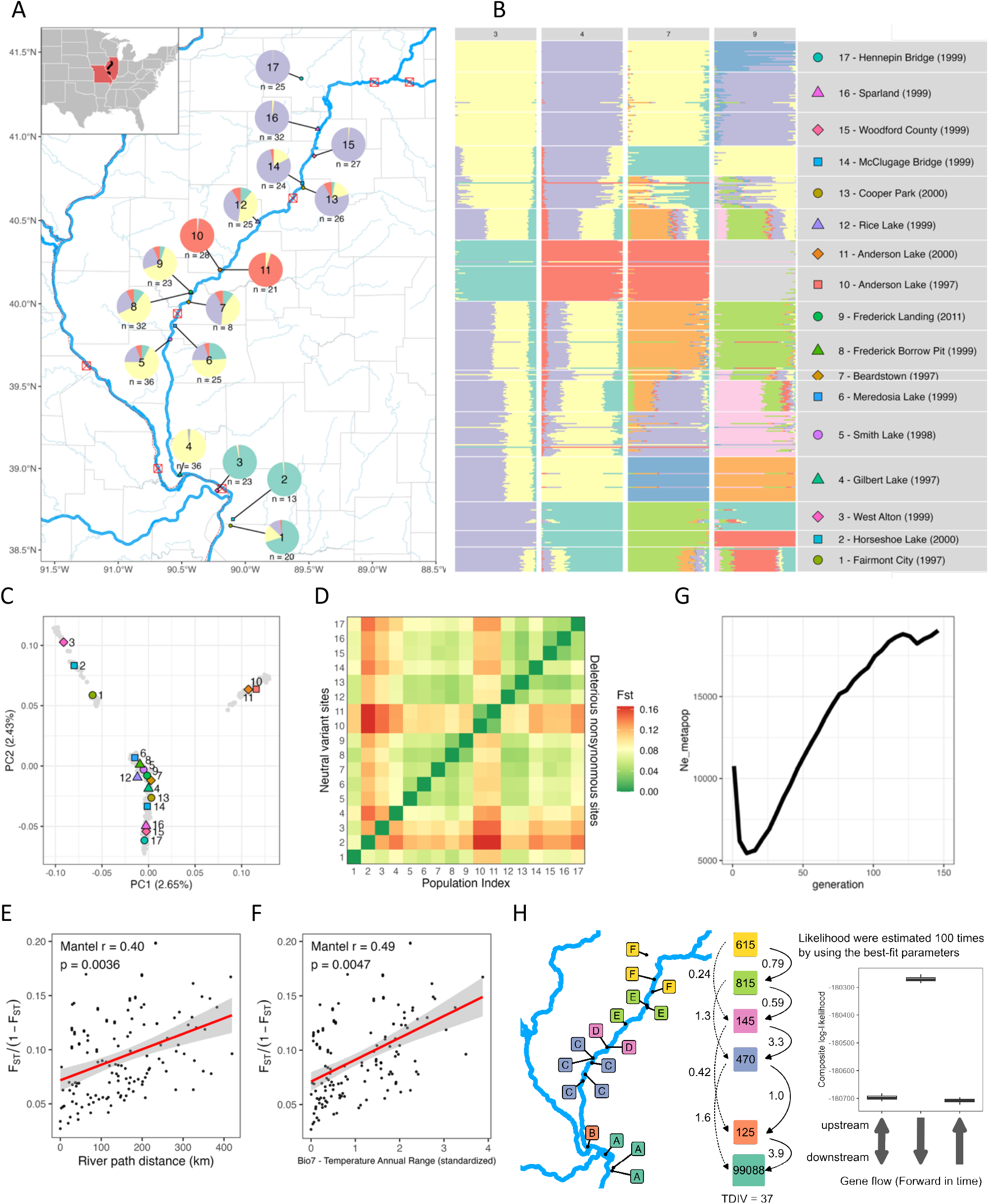
Population structure and isolation paFerns in *Boltonia decurrens*. **(A)** Map of *B. decurrens* populations, ordered and labeled from 1 to 17, from south to north. Pie charts at each site show ancestral genetic composition inferred from ADMIXTURE at K = 4. Red squares with crosses (⊠) indicate major dam and lock structures along the river system. (**B)** Individual ancestry proportions from ADMIXTURE. K = 9 yielded the lowest cross-validation (CV) error (see **Supplementary Fig. S5**). The legend on the right showing the combinations of symbol shape and color used for each population index (1–17) is applied consistently across all subsequent figures. **(C)** PCA of *B. decurrens*. Colored symbols are population medians and gray dots are individual values. The coordinates of the medians are manually adjusted to avoid overlapping. **(D)** Pairwise FST heatmap. The upper triangle shows pairwise F_ST_ estimated from all nuclear markers, and the lower triangle shows pairwise F_ST_ estimated from putatively deleterious nonsynonymous sites. **(E)** Isolation by river-path distance. **(F)** Isolation by environment – Temperature Annual Range (Bio7). **(G)** Recent demographic history, showing a pronounced bottleneck approximately 12–13 generations ago. **(H)** Recent demographic model of directional gene flow among *B. decurrens* populations under a stepping-stone framework.

### Intraspecific population genetic structure and gene flow

Genome-wide analyses revealed a clinal pattern in population structure across the 17 sampled *B. decurrens* populations, consistent with hydrochory as the primary dispersal mechanism (**Fig. 2A–D**). ADMIXTURE showed a clinal pattern of ancestry along a south–north axis (**Fig. 2B, Supplementary Fig. S5**). PCA similarly captured this gradient on PC 1 and 2 **(Fig. 2C**). However, populations #10 and #11 at Andersen Lake and the southernmost populations #1, #2, and #3 exhibited divergent ancestry from the otherwise clinal pattern (**Fig. 2A–B**). These divergent populations were not associated with the locks and dams along the Illinois River, suggesting that these types of river engineering were not a barrier to gene flow.

This clinal pattern in genetic structure is also consistent with analyses of isolation by distance and environment (**Fig. 2D**). Mantel tests showed that F_ST_ was significantly positively correlated with river path distance (Mantel’s *r* = 0.40, **Fig. 2E**), and was even more strongly correlated with Temperature Annual Range (Bio7) (Mantel’s *r* = 0.49; **Fig. 2F, Supplementary Fig. S6**), which showed the strongest association with genetic differentiation. The variation in Temperature Annual Range (Bio7) comes from the greater summer-winter temperature difference in the north relative to southern populations (**Supplementary Fig. S7**).

Next, we modeled contemporary gene flow using fastsimcoal2 to compare stepping-stone models in which populations were combined into six regional groups based on ADMIXTURE and PCA results (**Figs. 2B and 2C, respectively**), with recent gene flow modeled as either equally bidirectional, predominantly downstream, or predominantly upstream (**Fig. 2H**). Consistent with our hypothesis, the downstream-migration model had the highest likelihood (**Fig. 2H**). In addition, the Anderson Lake populations (#10 and #11; Group D in the model) were inferred to have a low effective population size and minimal incoming gene flow from upstream populations (Group E), consistent with restricted hydrological connectivity into the lake. The model also inferred a disproportionately large effective population size for the southernmost populations (#1–#3; Group A), nearly two orders of magnitude greater than that of other groups.

### Inbreeding, genetic diversity, genetic load

The population discontinuities in genetic structure in *B. decurrens* involving two geographic groups of populations also affected genetic diversity. Populations #10 and #11 (Anderson Lake) were genetic outliers characterized by the highest RoH, the lowest heterozygosity, and elevated Tajima’s D (**Fig. 3A, B**), together indicating inbreeding and recent population contraction. Despite Anderson Lake’s proximity to the main river channel, effective hydrological connectivity is likely currently limited. Downstream populations #1, #2, and #3— located below the confluence of the Illinois, Mississippi, and Missouri rivers—also showed distinct ancestry compositions despite receiving gene flow from upstream populations (**Fig. 2A– C, H**). In addition, this distinct composition did not show signatures expected under recent bottleneck or elevated inbreeding (**Fig. 3A, B**). The lack of genetic erosion in these distinctive populations suggests that the differentiation of #1–#3 is more consistent with ecological adaptation to environmental variation, such as more clay content in soil near the confluence (Smith et al., 1995), rather than a simple downstream colonization bottleneck.

**Figure 3.**
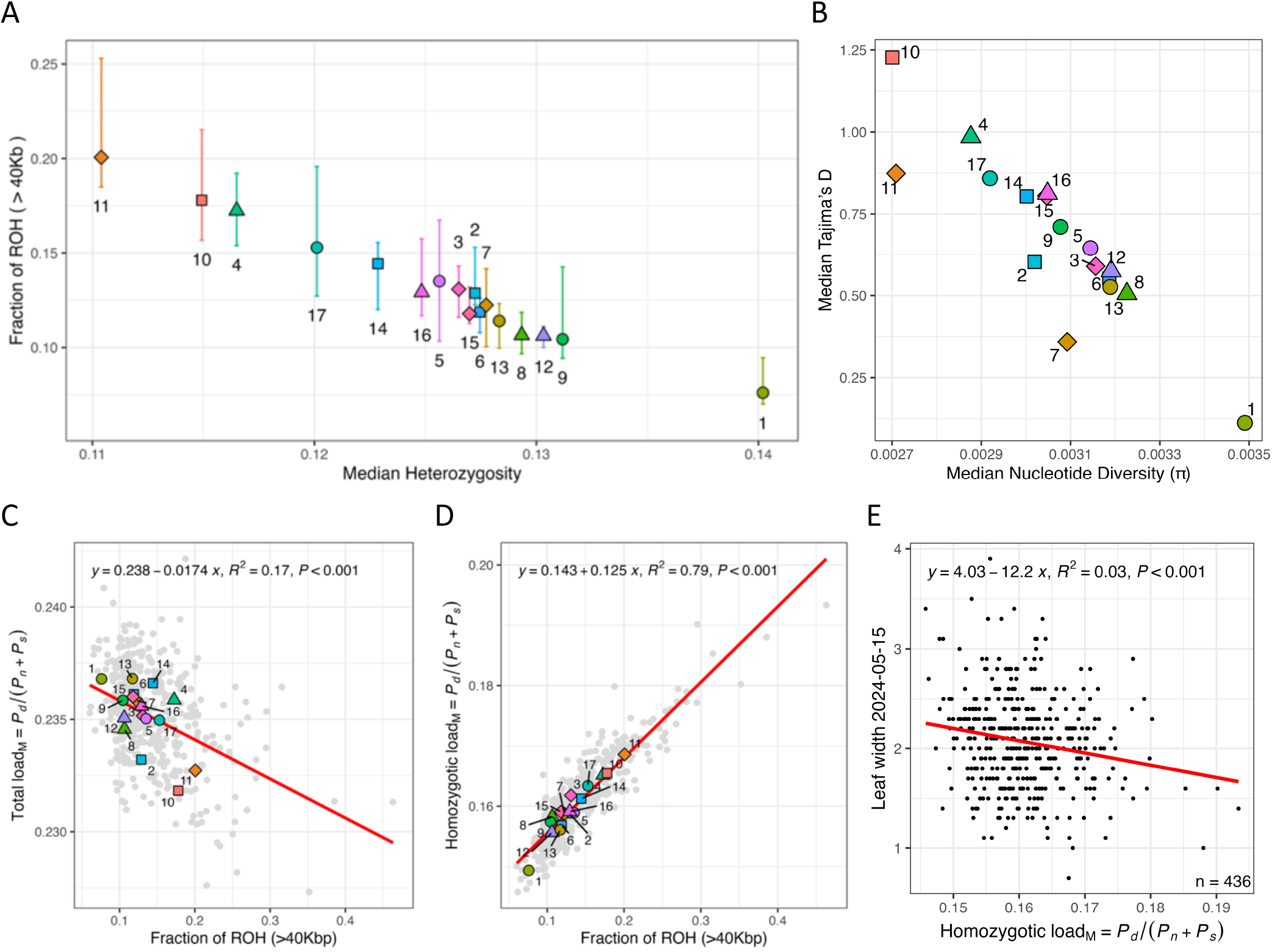
Population demography and genetic load in *Boltonia decurrens*. **(A)** Genome-wide runs of homozygosity (ROH) and heterozygosity across individuals. **(B)** Window-based paierns of genetic diversity and Tajima’s D estimated in 10 kb non-overlapping windows. **(C)** Total genetic load per individual, estimated from the cumulative burden of putatively deleterious nonsynonymous variants. **(D)** Homozygous genetic load (realized load), measured as the number of deleterious alleles in homozygous state per individual, reflecting the realized expression of deleterious mutations. **(E)** Association between homozygous genetic load and early vegetative growth (leaf width of the longest leaf on 2024-05-15).

In *B. decurrens*, analyses of genetic load varied by whether we examined total vs. homozygous (realized) genetic load. Across populations, total genetic load was negatively associated with inferred inbreeding; for example, populations #10 and #11 (Anderson Lake), which exhibited the highest RoH values (**Fig. 3A**), had the lowest total genetic load (**Fig. 3C**), whereas population #1 (Fairmont City), with the lowest RoH values (**Fig. 3A**), ranked among the highest for total genetic load (**Fig. 3C**). In marked contrast, homozygous load (i.e., realized load) was strongly positively associated with inbreeding as inferred from RoH (**Fig. 3D**), peaking in populations #10 and #11 and reaching its minimum in population #1. Together, these patterns suggest that although inbreeding may have purged the overall burden of segregating deleterious alleles in populations #10 and #11, these populations nonetheless carry the highest realized (homozygous) deleterious burden and therefore may be most susceptible to inbreeding depression. To evaluate whether deleterious variation affected plant fitness or performance, we tested associations between realized genetic load and all measured phenotypes (**Supplementary Fig. S8**). Realized genetic load was negatively associated with the width of the longest leaf at 2–2.5 months after planting—just prior to the earliest flowering (**Fig. 3E**; *h*^2^ = 0.739). Because leaf width at this stage is a proxy for juvenile leaf area and growth potential, this pattern suggests that realized genetic load may reduce performance during early vegetative development.

### The genetic basis of functional traits and climate adaptation

In the greenhouse, *B. decurrens* populations exhibited broad variation in vegetative and reproductive traits within each population. Life-history expression also varied among populations: 10–50% of individuals in each population flowered in their first year, while the remaining individuals flowered by their second year (**Fig. 4A**). All individuals that flowered in their first year also flowered in the second year. No source population showed a fixed life-history strategy. GWAS identified two adjacent loci associated with this life-history variation (**Fig. 4B, C, D**). Both peaks were located upstream of a gene (*SNRNP25*) encoding a U11/U12 small nuclear ribonucleoprotein, a core component of the minor spliceosome involved in removal of U12-type introns from pre-mRNA and previously linked to inflorescence development in *Arabidopsis* (Kim et al., 2010).

**Figure 4.**
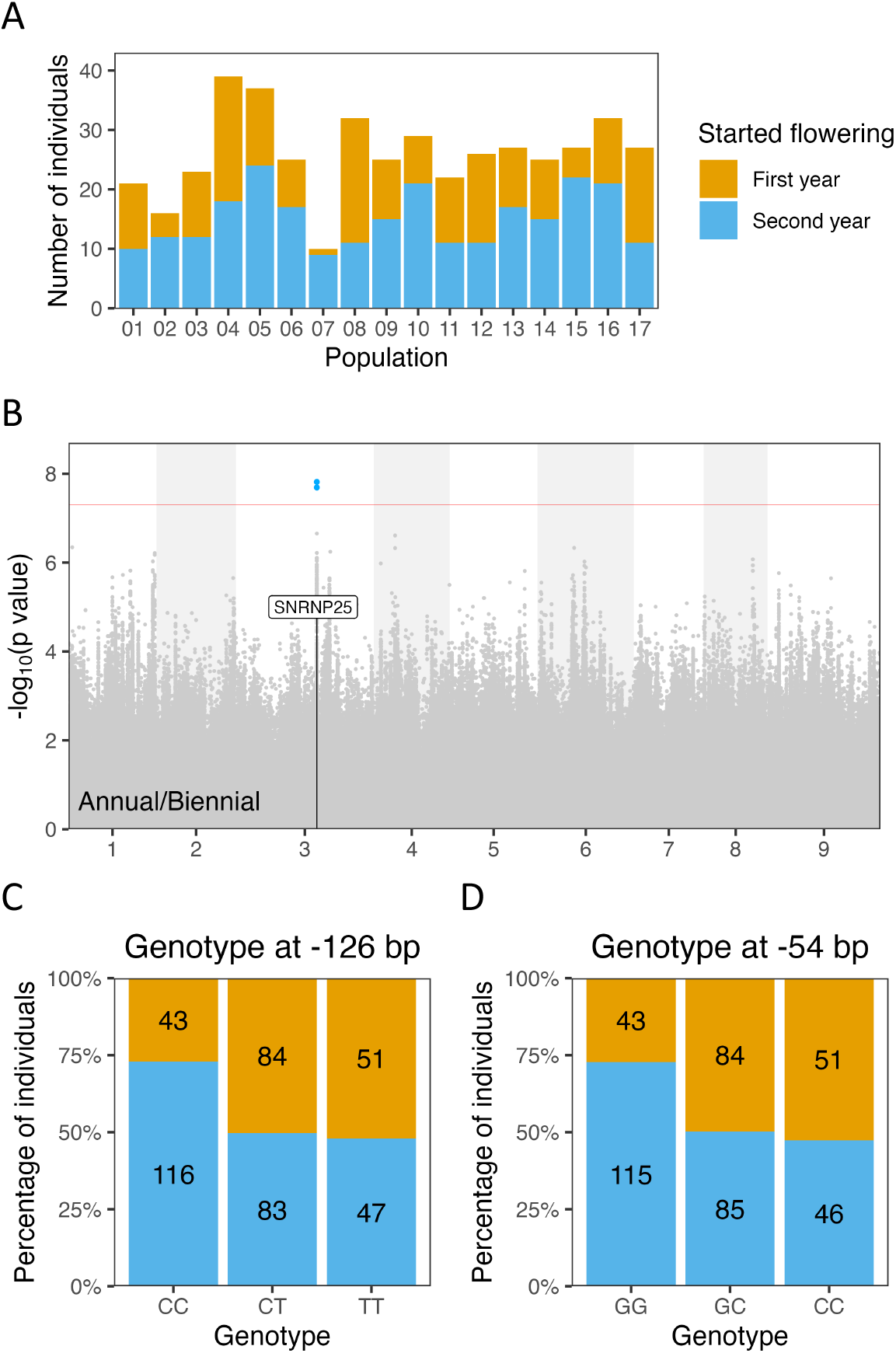
Variation in life history between first-and second-year flowering. **(A)** Life history variation among populations. All individuals that flowered in their first year also flowered in the second year. **(B)** GWAS results and candidate gene associated with life history. **(C)** Effect of genotype at the −126 bp site upstream of *SNRNP25*. **(D)** E?ect of genotype at the −54 bp site upstream of *SNRNP25*. Numbers indicate the number of individuals in each genotype class.

Both flowering time and stem length showed optimal ranges for the maximum reproductive output under greenhouse conditions. The plants started flowering around 100– 120 days after germination (**Fig. 5B**) and those with stem lengths of 50–60 cm (**Fig. 5E**) generated the greatest number of inflorescences in 2024. Flowering time and stem length were positively correlated (slope = 0.685, R^2^ = 0.36, *p* < 0.001). GWAS identified loci associated with flowering time (**Fig. 5C**; *h*^2^ = 0.964), stem length at flowering (**Fig.** 5F; *h*^2^ = 0.780), and number of stems when the first inflorescence matured (**Supplementary Fig. S10**; *h*^2^ = 0.256). Candidate genes near flowering-time association peaks included ***SKP1B***, a component of SCF ubiquitin-ligase complexes; ***ETO1***, a negative regulator of ethylene biosynthesis; ***CIPK5***, a calcium-sensor-interacting protein kinase; and ***FLX***, a regulator of FLC-dependent flowering (**Supplementary Table S3**). Candidate genes associated with stem length represented diverse growth-related processes, including putative receptor-kinase signaling (***At1g07650***), plastid isoprenoid biosynthesis (***DXS2***), ethylene-responsive transcription (***ERF003***), plasma-membrane proton transport (***AHA9***), small-RNA biogenesis (***DCL4***), and kinesin-related cellular functions (***KIN7K***) (**Supplementary Table S4**). Candidate genes associated with stem number included genes implicated in auxin signaling (***IAA13***), endosomal and vacuolar trafficking (***RAB7***), lipid hydrolysis and salt-stress responses (***LTL1***), and plant immune or temperature-dependent defense signaling (***RIN13*** and ***CHS3***) (**Supplementary Table S5**).

**Figure 5.**
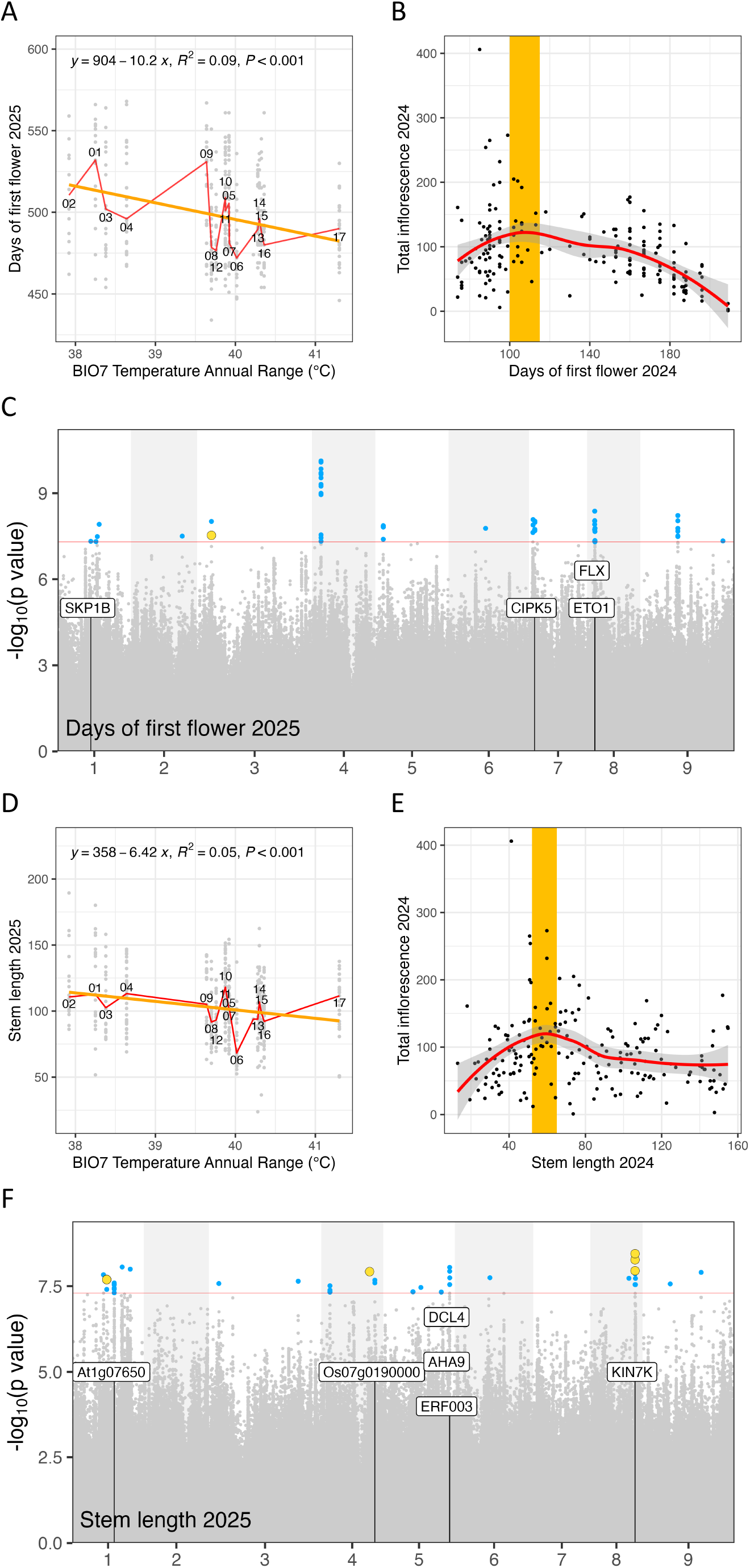
Functional trait variation and genetic associations. **(A)** Days to first flowering in 2025 showed a negative association with temperature annual range (BIO7). **(B)** Reproductive output in relation to days to first flowering in 2024. **(C)** GWAS results and candidate genes associated with days to first flowering in 2025. **(D)** Stem length in 2025 showed a negative association with temperature annual range (BIO7). **(E)** Reproductive output in relation to stem length in 2024. **(F)** GWAS results and candidate genes associated with stem length in 2025.

As described above, Temperature Annual Range (BIO7) showed the strongest association with genomic differentiation. We next asked whether this climatic gradient also corresponded to phenotypic and genomic signatures of local adaptation. The common garden showed clinal variation in adaptive traits in association with BIO7. With increasing BIO7, flowering time became earlier (**Fig. 5A**; slope = −10.2, *R*^2^ = 0.09, *p* < 0.001) and flowering stem length shortened (**Fig. 5D**; slope = −6.42, *R*^2^ = 0.05, *p* <0.001). Because these relationships were expressed under shared greenhouse conditions, they indicate that trait divergence along the BIO7 gradient is not solely attributable to environmental plasticity, but rather that earlier flowering and shorter plants originated in more northern populations.

We next tested whether *B. decurrens* exhibited genomic signatures of local adaptation across the environmental cline. We combined individual-based and population-based scans to leverage complementary assumptions about background structure (**Fig. 6**). At the individual level, we used PCadapt to identify loci with excess differentiation (**Fig. 6A**), and LFMM to identify variant sites associated with environmental variables while accounting for background structure (**Fig. 6B**). At the population level, we used BayPass to detect variant sites with elevated differentiation among populations (**Fig. 6C**) and variant sites associated with environmental variation (**Fig. 6D**). Across methods, these scans showed measurable concordance: 2,425 of 6,950,268 variant sites fell within the top 0.1% of test statistics in both PCadapt and LFMM, and 607 sites fell within the top 0.1% in both BayPass differentiation and BayPass environmental-association analyses. In total, 45 variant sites were consistently identified as outliers across all four scan types (**Fig. 6**, purple dots). To assess whether genomic signals of local adaptation also aligned with the genetics of adaptive traits, we next examined overlap between differentiation/environmental-association outliers and GWAS hits. Overlap was limited but non-random. LFMM shared four hits with loci identified by GWAS for stem length (**Fig. 6B**), and BayPass differentiation scans shared one hit with GWAS for time to flowering (**Fig. 6C**).

**Figure 6.**
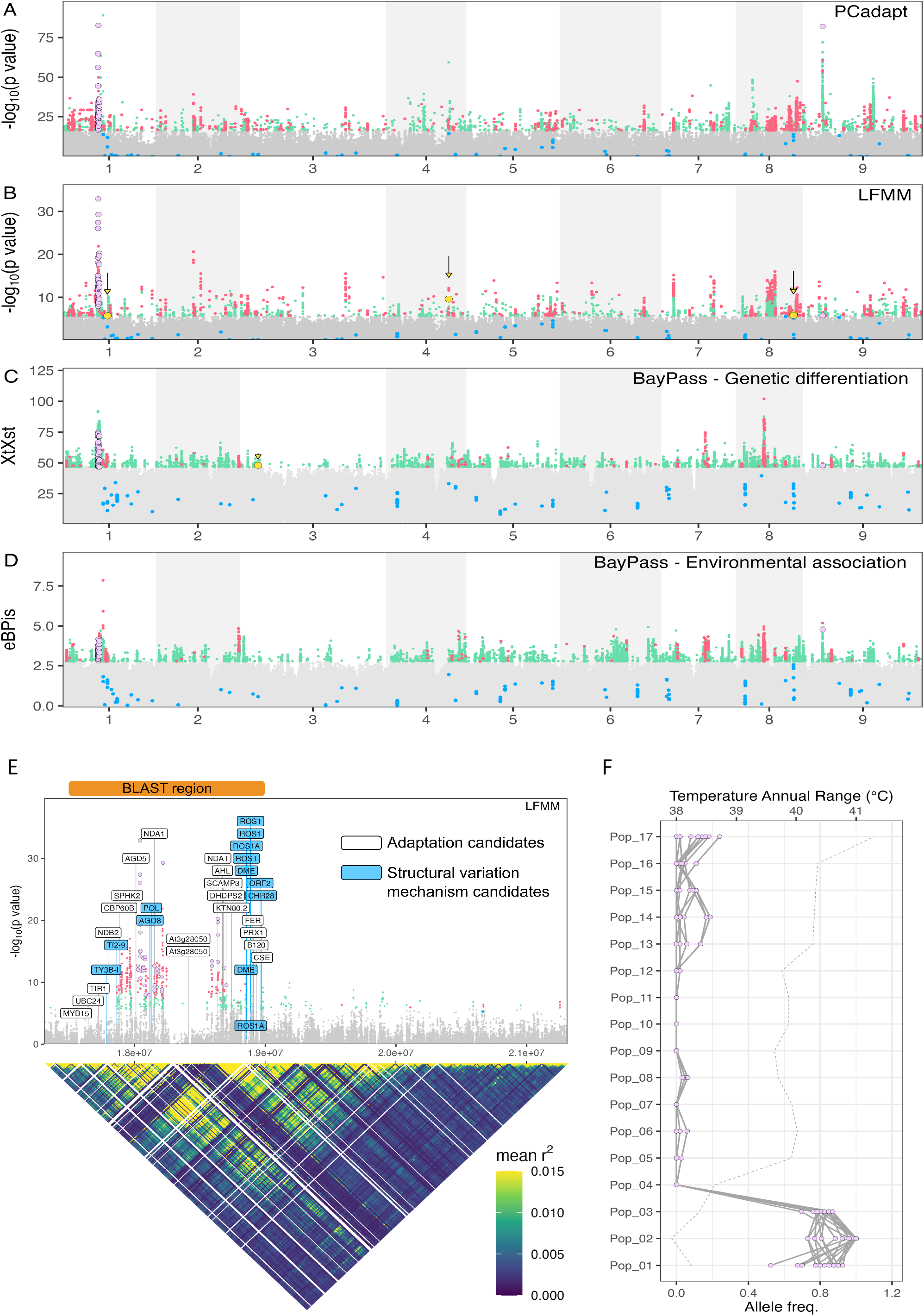
Genetic diFerentiation and genotype–environment association scans. **(A)** PCadapt outlier scan. **(B)** LFMM genotype–environment association scan; yellow points indicate the GWAS signal associated with stem length in 2025. **(C)** BayPass population-level genetic differentiation scan; yellow points indicate the GWAS signal associated with days to first flowering in 2025. **(D)** BayPass population-level environmental association scan. In **(A–D)**, blue points mark significant GWAS hits for days to first flowering, stem length, and stem number. Red points indicate SNPs jointly ranked in the top 0.1% within paired analyses: PCadapt ∩ LFMM or BayPass genetic differentiation ∩ BayPass environmental association. Purple points indicate variant sites detected by all four scan methods. **(E)** A haplotype block showing a structural variant and a potential hotspot for local adaptation. **(F)** Allele frequencies of variant sites detected by all four scan types, corresponding to the purple points in **(A–D)** on chromosome 1, across populations.

Across analyses, the strongest convergence among different approaches to identify adaptive loci occurred in a prominent region on chromosome 1 (**Fig. 6E**). Across 17.0–21.5 Mb on Chromosome 1, genome scans identified two dense clusters of positive hits flanking an interval with markedly reduced signal (**Fig. 6E**). Linkage disequilibrium summarized in 10-kb windows remained elevated across this region, consistent with a large, tightly linked haplotype block or putative structural polymorphism. The intervening low-signal interval may reflect reduced mapability to the reference genome, potentially caused by sequence divergence or breakpoint-associated rearrangement. Within the core 17.5–19.0 Mb interval, this block contains multiple genes with functions plausibly relevant to stress response, nutrient signaling, and growth regulation, including *MYB15, SPHK2, UBC24, TIR1, FER*, and *CSE*, together with a local enrichment of transposable-element and epigenetic-regulation machinery, including Ty3/Tf2/TNT1-like elements, *AGO8, DME, ROS1*, and *CHR28*. Allele frequencies at intersecting outlier sites across this block exhibited a pronounced association with BIO7, with strongly shifted haplotype frequencies in southern populations (#1–#3) relative to northern populations (#4–#17) (**Fig. 6F**). Thus, despite ongoing gene flow and metapopulation dynamics, signals of local adaptation in *B. decurrens* appear concentrated in a small number of regions, with the chromosome 1 haplotype block emerging as a candidate hotspot for ongoing and future adaptive differentiation along the climatic gradient.

## Discussion

In North America, up to 90% of floodplains have been converted for agriculture and other uses, and the remaining floodplain habitat has been functionally degraded (Tockner & Stanford, 2002); this floodplain degradation is closely tied to freshwater biodiversity declines. Against this backdrop, species conservation is entering a new phase in which reference-genome-enabled whole-genome resequencing, structural-variant discovery, and genotype-phenotype mapping can be combined with landscape genomics to generate mechanistic and management-ready insights for non-model species (Formenti et al., 2022; Theissinger et al., 2023). These approaches revealed that the threatened Illinois River floodplain endemic *B. decurrens* is closely related to *B. asteroides*, with genomic differentiation consistent with adaptation to a specialized floodplain niche. Interspecific gene flow was limited and predominantly directed from *B. decurrens* into *B. asteroides*, indicating that it is unlikely to constitute a major threat to the persistence of *B. decurrens*. Within *B. decurrens*, river-mediated connectivity and recent demographic history strongly shaped population structure, genetic diversity, inbreeding, and deleterious genetic load, highlighting the importance of maintaining connectivity among populations. Common-garden analyses further revealed substantial within-population variation in flowering time (first year vs. second year; days of the year), and height of flowering stems, together with genomic associations suggesting that these traits have been shaped by heterogeneous selection under altered flood regimes. Finally, landscape-genomic and genome-wide association analyses identified candidate loci associated with environmental and phenotypic variation, and indicated that structural variants may constitute important mechanisms underlying adaptation, despite ongoing gene flow and metapopulation dynamics. Collectively, these findings support a conservation strategy that maintains population connectivity and genetic health while accounting for local adaptation in floodplain endemics.

### Evolutionary origin and species boundaries

Accurately defining the evolutionary unit to be protected and determining whether interspecific hybridization threatens genetic integrity are fundamental prerequisites for species conservation (Fraser & Bernatchez, 2001; Grabenstein & Taylor, 2018; Rhymer & Simberloff, 1996). Phylogeny reconstruction placed *B. decurrens* as a monophyletic lineage sister to a paraphyletic assemblage of *B. asteroides* and *B. diffusa*, supporting the genetic distinctiveness of *B. decurrens* while also indicating that additional taxonomic work is needed to clarify species boundaries between *B. asteroides* and *B. diffusa*. Notably, our *B. asteroides* samples were collected from central North America and therefore fall well outside the geographic range of *B. asteroides* sensu stricto, which is restricted to the vicinity of the Susquehanna River in Pennsylvania and adjacent Maryland (Nesom, 2026). We also detected interspecific hybridization between *B. decurrens* and *B. asteroides* south of *B. decurrens*’ main geographic range. Hybrids primarily reflected gene flow from *B. decurrens* into *B. asteroides* (**Supplementary Fig. S3B**), contrary to the expectation that the more widespread species would be the predominant donor because of its larger population size. Although one possibility is that retention of *B. decurrens* alleles in these hybrids confers fitness advantages in more southern floodplain environments, hybrids are rare and have not been documented to form stable populations, potentially because of reduced hybrid fitness; however, the relative fitness of hybrids vs. parental species needs additional study to test this hypothesis. Moreover, all hybrid samples were from herbarium collections, which may bias towards morphologically unusual individuals that appear especially noteworthy to collectors (Wu et al., 2023) but may not form persistent populations. Taken together, these results indicate that *B. decurren*s is genetically distinct from its closest relatives and that interspecific hybridization and introgression does not appear to represent a threat to its genetic integrity, as was found previously using allozyme markers (DeWoody et al., 2011).

Results of the present study also suggest that *B. decurrens* might have evolved via geographical and ecological speciation, budding from the widespread *B. asteroides*. Both the sister relationships between the two species and the stark contrast between the narrow floodplain-restricted range *of B. decurrens* and the broad distribution of *B. asteroides* are consistent with a progenitor–derivative (budding) speciation scenario, where a small, geographically marginal lineage became isolated and differentiated from a widespread progenitor (Crawford, 2010). Furthermore, the southernmost population of *B. decurrens* (population #1) harbored the highest genetic diversity and effective population size; this population may represent the origin and primary reservoir of diversity for *B. decurrens*, with subsequent northward expansion generating founder effects that reduced diversity in northern populations. This pattern may have been reinforced by predominantly downstream gene flow, which could concentrate genetic variation in southern downstream populations and facilitate recolonization following catastrophic floods (Fink et al., 2022; Love, Maggs, Murray, & Provan, 2013). Although this pattern could also be the product of interspecific gene flow and introgression with *B. asteroides*, our results indicate that the primary direction of introgression was from *B. decurrens* into *B. asteroides* rather than the reverse. We did not detect excessive allele sharing between population #1 and *B. asteroides* relative to other *B. decurrens* populations (#2–#17), arguing against substantial recent introgression as the primary cause of elevated diversity (**Supplementary Fig. S3A**). This interpretation is also consistent with postglacial phylogeographic patterns reported for eastern North American plants, in which southern refugial populations retained greater diversity and subsequently expanded their ranges northward into areas opened by Laurentide ice-sheet retreat (Dalton et al., 2023; Mohn et al., 2021; Park & Donoghue, 2019). Taken together, these lines of evidence support a speciation scenario in which *B. decurrens* likely originated near the southern margin of its present range, expanded northward via flood-caused backflows, and accumulated a latitudinal diversity gradient shaped by founder effects and directional seed dispersal, while experiencing limited contact with *B. asteroides* outside its contemporary core distribution.

### Demographic changes, gene flow, and genetic diversity in *B. decurrens*

In *B. decurrens*, multiple genomic lines of evidence argue that rarity is coupled with **recent demographic instability**. Demographic simulations supported a severe recent bottleneck followed by a population rebound (**Fig. 2G**). Furthermore, across the genome, *B. decurrens* exhibited consistently lower nucleotide diversity and more positive Tajima’s *D* values than its widespread congener, *B. asteroides* (**Fig. 1C**). Together, these patterns suggest that recent population declines and habitat fragmentation have increased both inbreeding (**Fig. 3A**) and realized genetic load (**Fig. 3D**), pointing to elevated vulnerability.

In the nuclear genome, we detected a broad **clinal pattern in population structure** along the river corridor, consistent with isolation-by-distance (IBD) (**Fig. 2B**). The finding indicates that the populations are connected, but gene flow has occurred across regional rather than panmictic scales. This contrasts with earlier allozyme work reporting significant differentiation but no isolation-by-distance (IBD) (Dewoody et al., 2004). One possibility for the difference in the results of the two studies is the timeframe of when populations were sampled relative to the major demographic decline and subsequent population re-expansion detected in the present study. Dewoody et al. (2004) sampled between 1984-1989, potentially before the large demographic recovery, whereas the current study sampled between 1997-2011, after the 1993 and 1995 floods that resulted in population rebounds (Smith et al., 1998).

Plastome patterns further support a recent demographic contraction followed by expansion, while also providing direct evidence for the capacity of seeds to move long distances. Most populations share a single ancestral plastome haplotype, with multiple low-frequency satellite haplotypes derived from it, a configuration consistent with a bottleneck and subsequent demographic rebound (**Fig. 1D**). The widespread sharing of the ancestral haplotype provides robust evidence that extreme flooding can generate backflow-mediated upstream dispersal, moving seeds from downstream sources into upstream reaches (Smith et al., 1998; Smith & Keevin, 1998; Smith & Moss, 1998).

### Adaptive traits and genetics

In the current study, despite a recent demographic bottleneck, we recorded substantial variation in adaptive traits within and among populations of *B. decurrens*, including flowering time, flowering stem length, and number of flowering stems. In the common garden, which provided a relatively stable and homogeneous environment, fitness peaked around a single intermediate optimum of flowering time (**Fig. 5B, E**). This pattern reinforces the idea that the broader trait variation observed within and across populations has been maintained by heterogeneous selection in nature.

Another trait that varied within populations of *B. decurrens* was their year of first flowering. Previously, plants were reported to have annual, biennial, or perennial lifespans (Smith et al., 2005), but our common garden experiment showed that all plants are perennials but differ in whether they first flowered in their first or second years. Flowering in the first year was previously observed to be dominant in populations that experienced natural flow regimes with annual spring floods (Smith et al., 2005), yet it never reached above 50% of any sourced populations in the common garden. The altered flood regimes experienced by *B. decurrens* in recent years, with their unpredictable timing and extreme magnitudes, may have reshaped this trait, from one that dominantly flowered in the first year (previously reported as an annual life history) to one that shows more balance among those that first flower in their first and second years, which may serve as a hedge against unpredictable environmental conditions. This rapid evolution in this trait may be possible because it is associated with only two mutations (a single locus) upstream from a pre-mRNA processing gene, which has been reported to control inflorescence development (Kim et al., 2010). This shift in life-history could occur rapidly if it is under strong balancing selection because it is controlled by a single genetic locus.

### Local adaptation and structural variants

Given the phenotypic clines observed in flowering time and stem length, we tested whether *B. decurrens* nonetheless retains detectable genetic signatures of local adaptation despite the demographic and ecological conditions that may constrain it. A persistent limitation in plant conservation genomics is that signatures of local adaptation are often inferred from landscape-genomic patterns alone, without corresponding phenotypic evidence. As a result, it is frequently unclear how much of the observed genomic differentiation reflects functionally important trait variation. Our dataset offers an unusual opportunity to address this gap by integrating landscape genomics with GWAS for functional traits. The concordance between loci associated with flowering time and stem length and those identified by differentiation and genotype–environment association analyses indicated that some loci involved in local adaptation have a demonstrable functional basis (**Fig. 6A–D**). The non-overlapping candidate loci identified by the different landscape-genomic scan approaches may indicate additional selective axes not captured in our common garden, including hydrologic regime, nutrient conditions, sediment dynamics, or biotic interactions. Integrating these complementary approaches therefore provided a more discriminating view of local adaptation: not all genomic outliers can yet be assigned a phenotype, but some could now be linked to adaptive trait variation with much greater confidence.

Our results provide a compelling example of how linked loci may form a hotspot for local adaptation despite ongoing gene flow and metapopulation dynamics. The strongly differentiated block spanning >0.5 Mb on chromosome 1 is consistent with a putative structural variant that may facilitate divergence by maintaining sets of co-adapted alleles in different portions of the river corridor. Structural variants—including inversions, duplications, insertions, and deletions—are increasingly recognized as important substrates for adaptation because they can generate large functional effects through a single mutational event and preserve favorable allelic combinations despite ongoing gene flow (P. Battlay et al., 2025; Paul Battlay et al., 2025; Kuo, Wright, Small, & Olsen, 2024). In *B. decurrens*, the distinctiveness of this genomic block between southern populations (#1–#3) and northern populations (#4–#17) is particularly associated with the divergence in annual temperature range (BIO7) and the ecological transition associated with the confluence of major rivers. From a conservation perspective, these results suggest that southern and northern populations may warrant consideration as two evolutionarily significant units, with management aimed at preserving adaptive variation in both parts of the species’ range. More broadly, this result illustrates how structural variation can provide a plausible mechanistic basis for adaptive divergence even in a threatened floodplain endemic shaped by severe drift, high gene flow, and demographic instability.

### Recommendations for conservation management

Despite a strong recent bottleneck, *B. decurrens* retains substantial within- and among-population adaptive trait variation, even in the most isolated and inbred Anderson Lake populations (#10–11), where plants still exhibited comparable trait variance. Thus, we recommend that conservation efforts prioritize *in situ* conservation to preserve diversity in *B. decurrens* while multiple wild populations with meaningful variation persist. The maintenance of variation within populations of *B. decurrens* appears tightly coupled to hydrologic processes; we therefore recommend prioritizing management to restore floodplain processes such as spring flooding that sustains recruitment pulses (e.g., flood backflow) and population connectivity (Murray et al., 2019; Sander et al., 2018).

We would also recommend prioritizing management for the most diverse populations in the southern part of the species range, especially Fairmont City (#1), and the least connected populations at Anderson Lake #10–11. The Fairmont City population (#1) shows the highest and relatively stable effective population size, the lowest inbreeding, and the lowest realized genetic load. Notably, this population occurs on land that was formerly a horseshoe lake, but is now adjacent to Interstate I-55, has been levied to prevent flooding, and is surrounded by heavy development. Ideally, we recommend river engineering efforts to restore natural flood regimes and genetic connectivity. Barring that, we recommend management to maintain the open, early successional conditions preferred by *B. decurrens*. Similarly, we recommend restoration of floodplain connectivity in the most isolated populations at Anderson Lake. If near-term restoration of floodplain connectivity is infeasible, targeted population augmentations could be considered for these highly isolated populations However, any augmentation should be carefully considered, given the risks of increasing maladaptation or introducing deleterious alleles (Aguilar-Gómez et al., 2025). In practice, this argues for prioritizing hydrologic and habitat restoration first and using assisted movement only when isolation and demographic risks are high, and donor choice is justified by genetic, ecological, and trait-based evidence.

## Acknowledgements

We thank Kenneth Olsen for hosting W.-H. K and E. A.-M. at Washington University in St. Louis and providing computational resources for the analyses in this study. We thank Paige Mettler-Cherry, Jenny Mullikin, and Christopher Sedivy in National Great Rivers Research & Education Center for providing suggestions and discussion of the ongoing *B. decurrens* conservation efforts. We thank Taylor Gray, Largo Haight, and Julie Marik for assistance with managing plants in the Oertli Hardy Plant Nursery. We thank Jillian Cieslik for providing instructions on running fastsimcoal2 models. We thank Cooper Call, Jillian Cieslik, Sarah Herzog, Binoshi Hettihewa, Alexander Linan, Brock Mashburn, Jonas Mendez Reneau, Kenneth Olsen, Jordyn Riehn, and Akiva Zeff for providing comments and edits to the manuscript. This study was funded by the Institute of Museum and Library Services Grant MG-253252-OMS-23.

## Statements of ethics and integrity policies

### Data availability statement

All raw sequencing data are available under NCBI BioProject: PRJNA1217859 (reviewer link: https://dataview.ncbi.nlm.nih.gov/object/PRJNA1217859?reviewer=egfr8l9mef085tt68tqpinlt1a). The reference genome is available under NCBI BioProject: PRJNA1456120 (hap1) and PRJNA1456119 (hap2). All scripts to generate the primary figures are deposited at GitHub (https://github.com/kuowenhsi/MOBOT_Boltonia). All processed sequencing data, including VCF files, are archived at DRYAD (10.5061/dryad.44j0zpcwn).

### Ethics approval statement

No ethics approval was required for this study because the research involved plants and did not involve human participants or vertebrate animals.

## Author Contributions

W.-H. K conducted the experiments, analyzed the data, and wrote the manuscript. E. A.-M. conducted the gene flow analyses and wrote the manuscript. B. B. conducted DNA extractions. E. Co. conducted the experiments and analyzed the preliminary data. M.E. curated and maintained seed accessions, retrieved seed samples for experimentation, field collected *B. diffusa* leaf material, minor revisions to manuscript. J. H. and M. M. J. conducted the experiments and made minor revisions to the manuscript. R.S. curated the seed accession data and made minor revisions to the manuscript. B.D., N.D., S. W., and M. W. conducted the phenotyping experiments. C. E. Ed. and M.A.A. conceived the project, obtained funding for the project, conducted the experiments, and edited the manuscript.

## Tables and Figures

**Supplementary Fig. S1.**
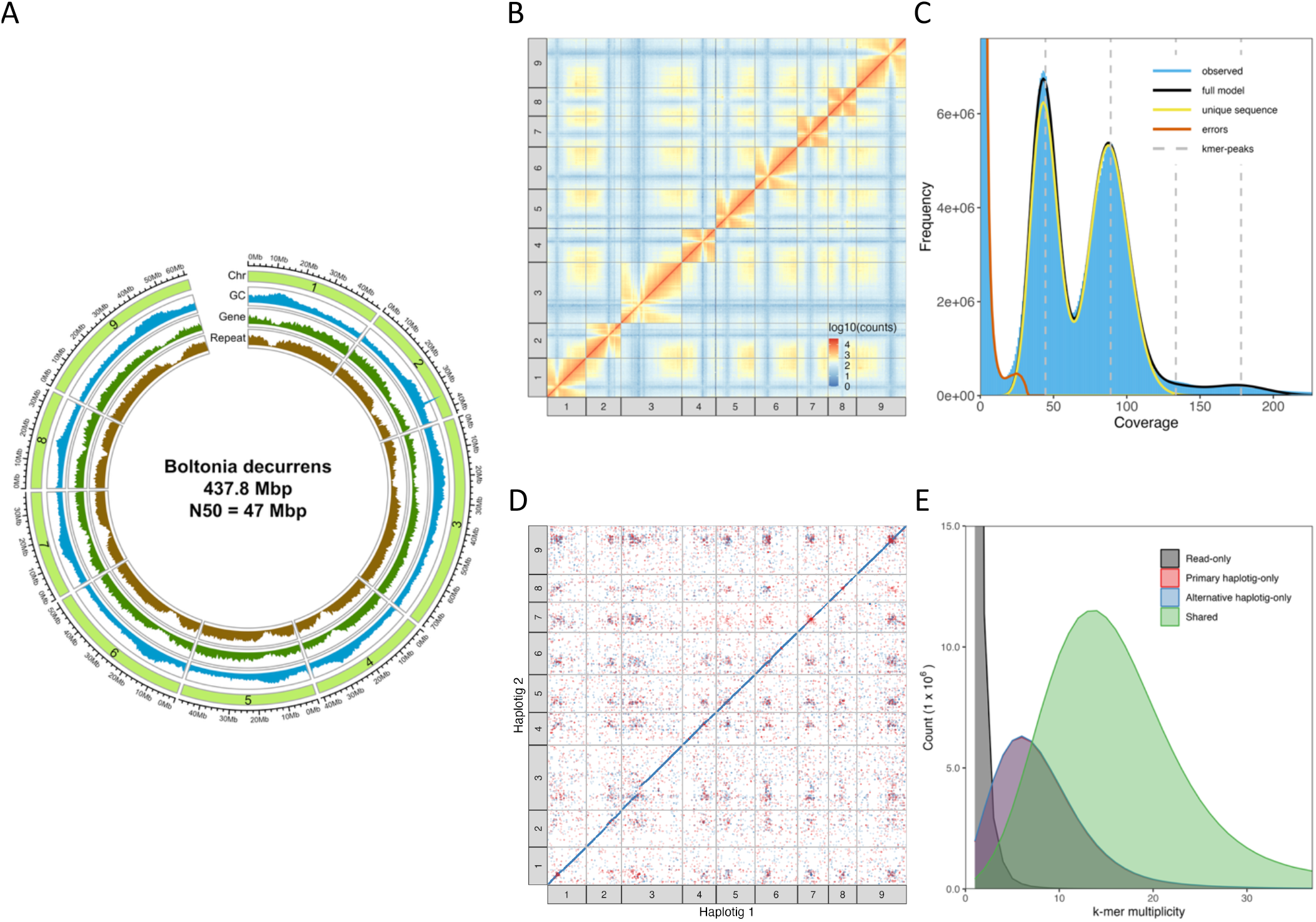
Genome assembly statistics for *Boltonia decurrens*. **(A)** Circular genome plot summarizing chromosome-scale features of the *B. decurrens* assembly, including GC content, gene density, and repetitive elements across scaffolds. **(B)** Hi-C contact heatmap showing chromosome-scale proximity interactions, with strong signal along the diagonal indicating accurate scaffolding and well-resolved chromosome structure. **(C)** GenomeScope k-mer profile based on PacBio HiFi reads (k = 21), estimating a haploid genome size of ~431 Mb with moderate heterozygosity (~1.6% homologous genome copies differ at a nucleotide position), and a low sequencing error rate (~0.15), with a high model fit (0.94). **(D)** Whole-genome alignment between the primary (haplotig 1) and alternate (haplotig 2) assemblies, illustrating high synteny and collinearity between haplotypes. **(E)** Merqury k-mer– based assembly quality assessment. K-mer completeness is ~81–82% for each haplotype individually and 99.3% when both haplotypes are combined, indicating nearly all read k-mers are represented in the phased assembly. Consensus quality scores (QV ≈ 38.6–38.7) correspond to an estimated base-level error rate of ~1.3–1.4 × 10^−4^, consistent with a highly accurate, chromosome-scale reference genome for *B. decurrens*.

**Supplementary Fig. S2.**
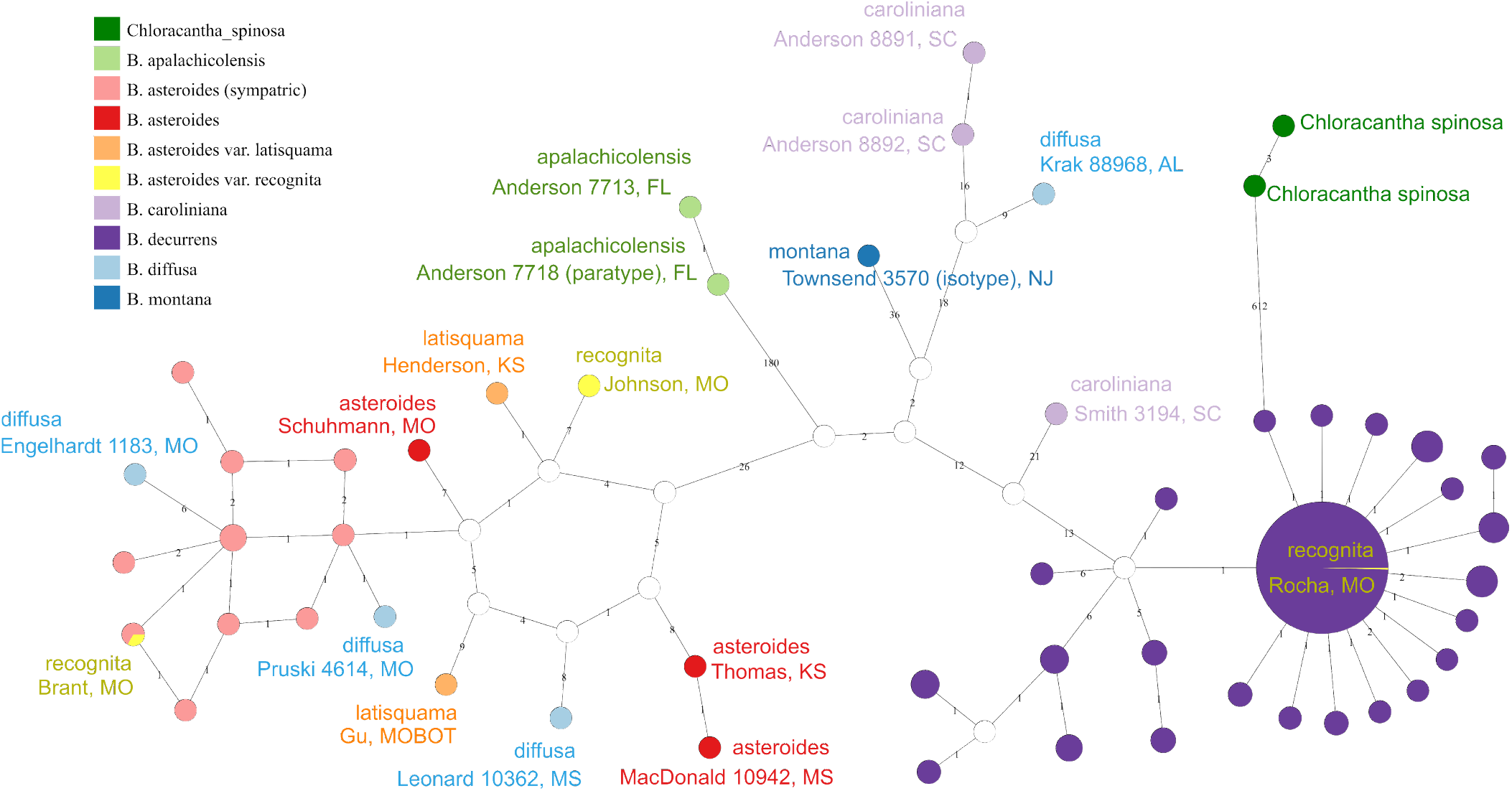
Plastome haplotype network. The specimen is named by [collector + collection number], [collection state code]. See **Supplementary File 1** for detailed information.

**Supplementary Fig. S3.**
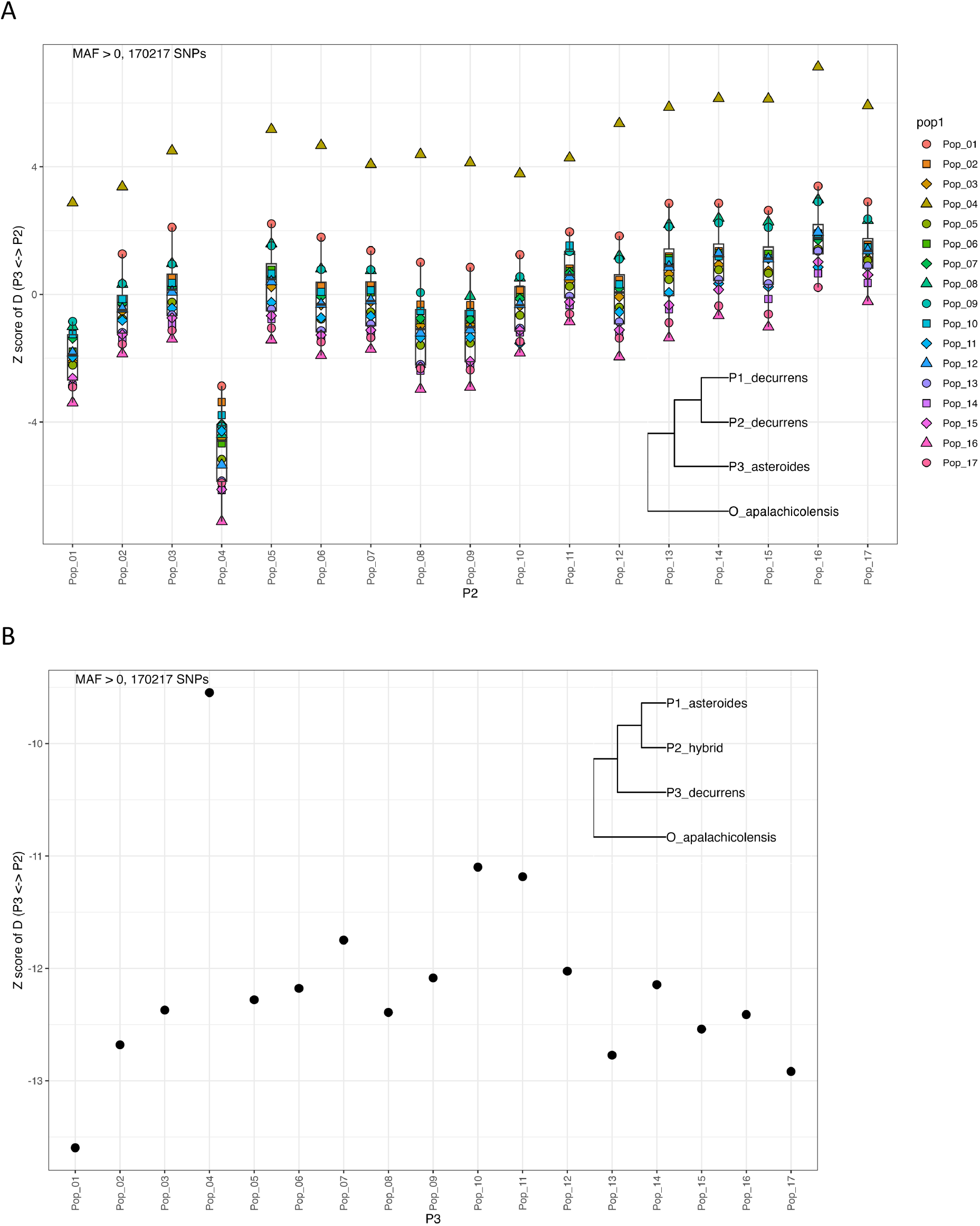
*D*-statistics testing asymmetrical sharing of derived allele among *D* (P1, P2, P3, Outgroup). **(A)**, *D*-statistics to test whether there is a certain *B. decurrens* population received more gene flow (derived allele) from *B. asteroides* than other *B. decurrens* population. The statistic result shows that *B. decurrens* population 04 received more gene flow from *B. asteroides*. However, it is not consistent with the ADMIXTURE results, nor the PCA result. We suggest that any further inference requires additional investigations. In the current study, we conservatively state that *B. decurrens* population #01, although contains shared genetic composition with *B. asteroides* (**Fig. 1A**), did not have higher gene flow with *B. asteroides* than any other *B. decurrens* populations. **(B)**, D-statistics to test whether there is a certain *B. decurrens* population contributing more to the hybrids, which grew outside the southern boundary of *B. decurrens*. We found that all *B. decurrens* populations had significant amount of shared derived alleles with the hybrids (*z* < −3). Among which, the *B. decurrens* population #01 has the lowest *z* value. It supports the hypothesis that the southern *B. decurrens* populations contributed more to the formation of the hybrids. **Note:** *D*-statistics were calculated in ADMIXTOOLS, which uses *D* = (BABA − ABBA) / (BABA + ABBA); therefore, the sign is reversed relative to some presentations of the ABBA–BABA test. *z*-scores were calculated as *z* = D / SE using a block jackknife, and |*z*| > > was considered evidence of significant deviation from zero. Under the ADMIXTOOLS convention, positive *z*-scores indicate excess allele sharing between P1 and P3, whereas negative *z*-scores indicate excess allele sharing between P2 and P3.

**Supplementary Fig. S4.**
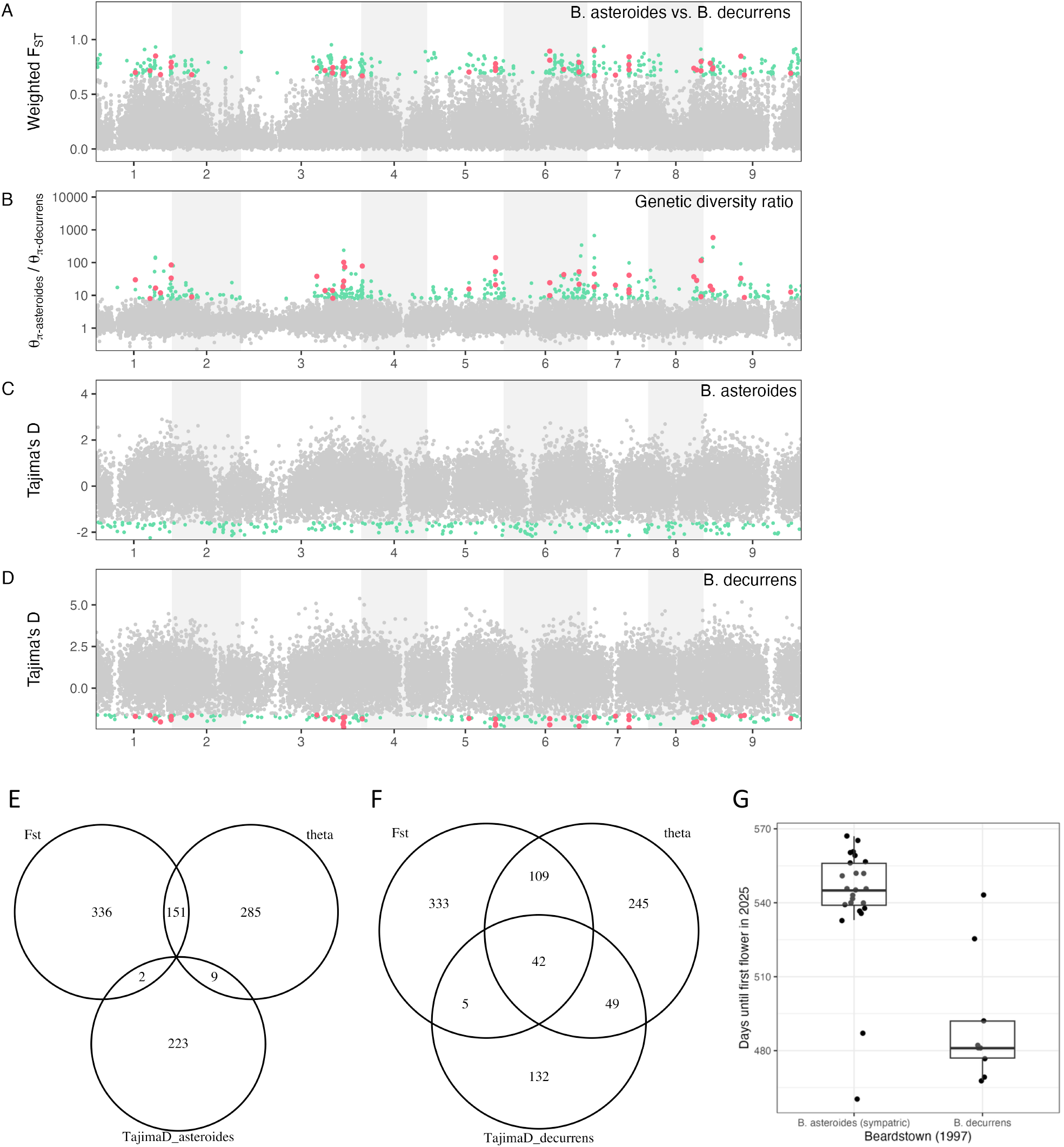
Genomic scans for positive selection in *Boltonia decurrens* relative to *B. asteroides*. **(A)** Pairwise FST between *B. decurrens* and *B. asteroides* across the genome, calculated in 10 kb non-overlapping windows. **(B)** Ratio of window-based genetic diversity (*π*) between *B. decurrens* and *B. asteroides*, calculated in 10 kb non-overlapping windows. **(C)** Tajima’s D estimated within *B. asteroides* using 10 kb sliding windows with a 5 kb step. **(D)** Tajima’s D estimated within *B. decurrens* using 10 kb sliding windows with a 5 kb step. In panels A–D, green points mark windows in the top 0.1% of each genome-wide distribution, and red points highlight windows that fall in the top 0.1% of all three scans: genetic diversity ratio, F_ST_, and Tajima’s D within *B. decurrens*. **(E)** Venn diagram showing the overlap in outlier variant sites (top 0.1% of each distribution) identified by scans of genetic diversity ratio, F_ST_, and Tajima’s D for *B. asteroides*. **(F)** Venn diagram showing the overlap in outlier variant sites identified by scans of genetic diversity ratio, F_ST_, and Tajima’s D for *B. decurrens*. **(G)** Comparison of days to first flower in 2025 between *B. asteroides* and *B. decurrens* individuals collected from the same locality at Beardstown, Illinois (Population #07). *B. asteroides* exhibits substantially delayed flowering relative to *B. decurrens* under common greenhouse conditions.

**Supplementary Fig. S5.**
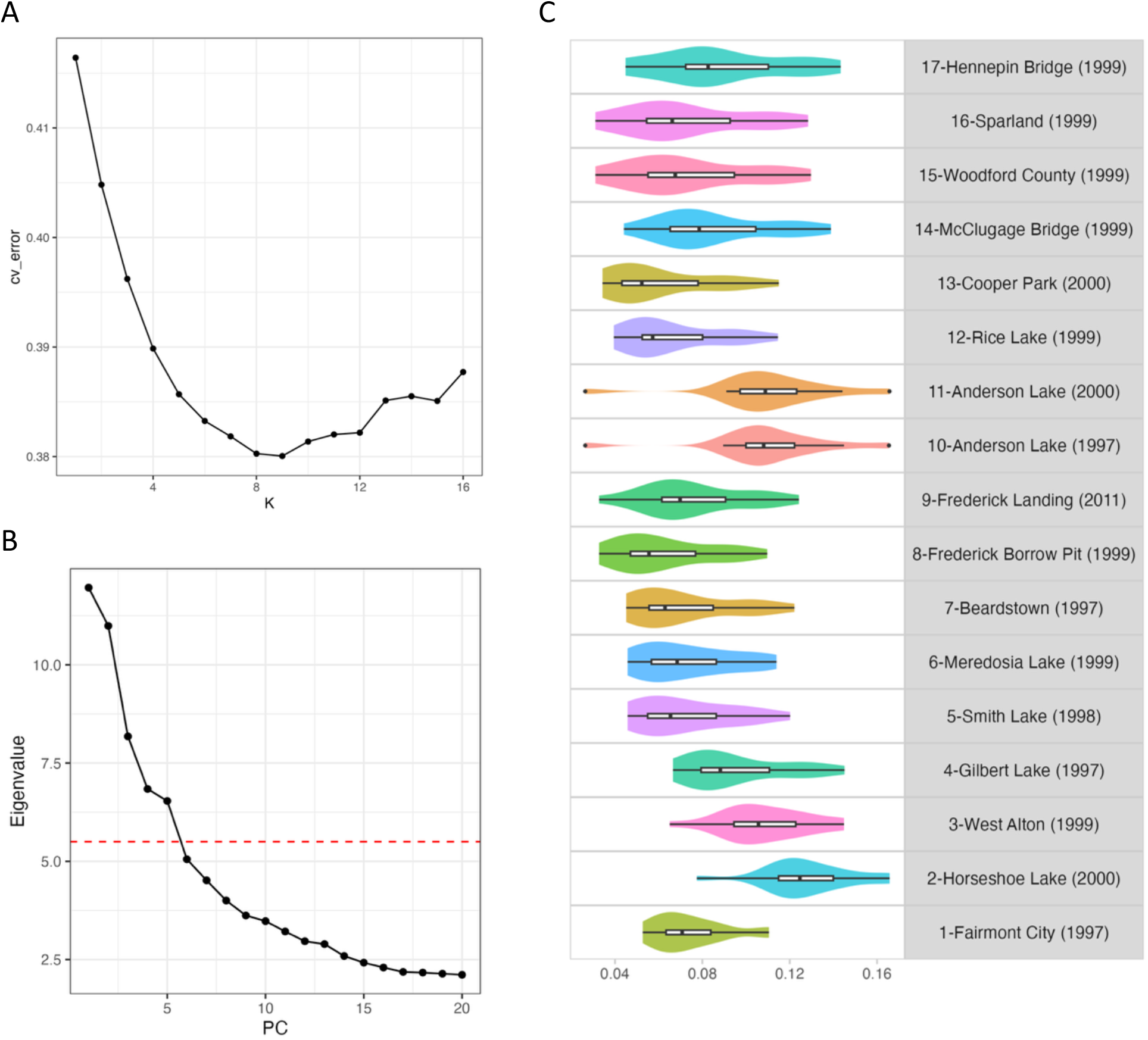
Additional information for population structure analyses. **(A)** Cross-validation (CV) error from ADMIXTURE runs across values of *K*. The minimal CV error occurs at K = 9, which was used as the primary clustering solution in the main analyses. **(B)** Scree plot showing the eigenvalues of principal components from the PCA. The first five PCs were retained and used both to set the dimensionality of the latent factor matrix in LFMM and to define the K values used in PCadapt. **(C)** Pairwise F_ST_ values summarized by focal population, illustrating how the strength of genetic differentiation varies among *B. decurrens* populations.

**Supplementary Fig. S6.**
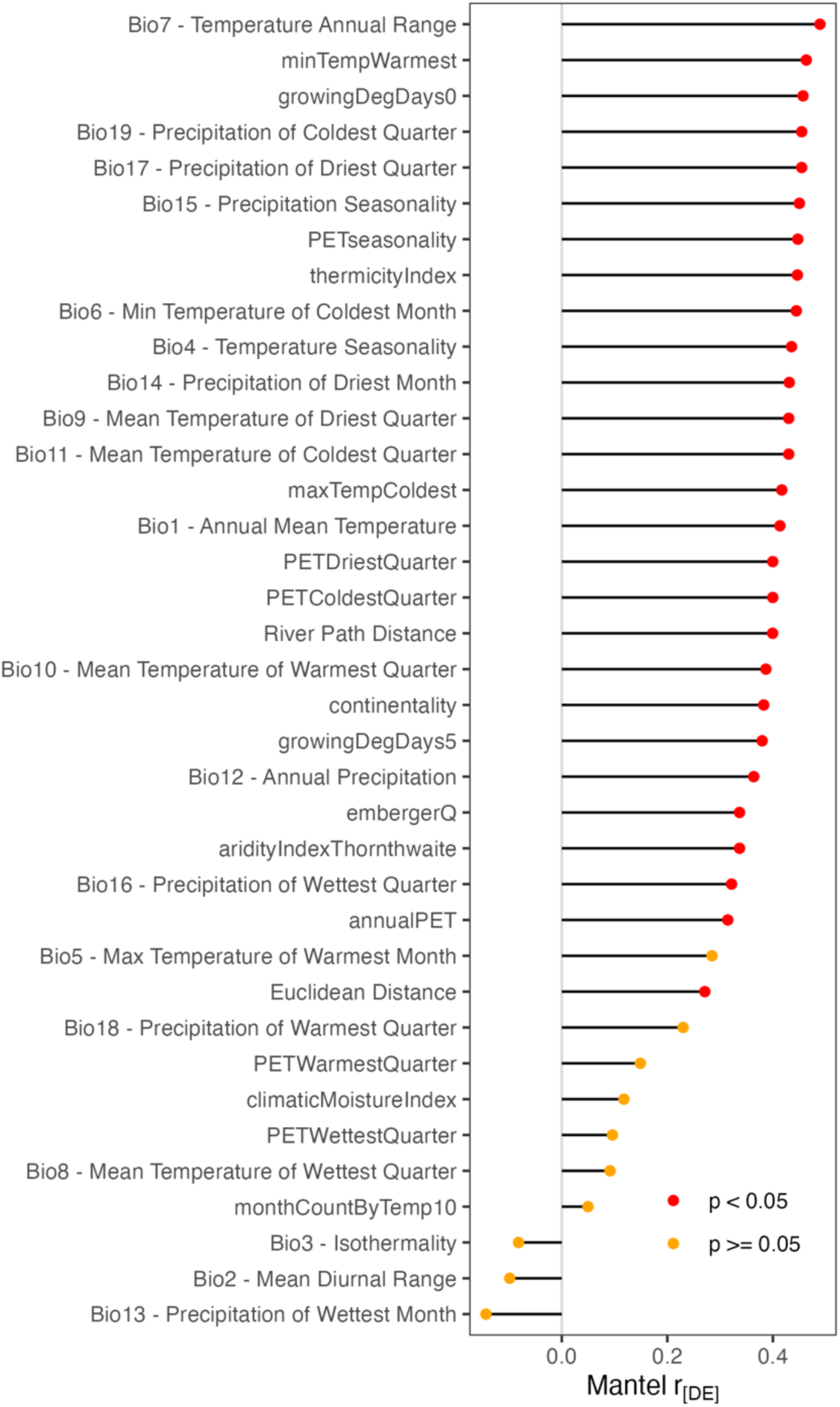
Mantel correlations for isolation by environment. Mantel correlation coefficients (*r*) between pairwise genetic differentiation and pairwise environmental distances across multiple predictors, including 19 bioclimatic variables from the WorldClim2 database (Fick & Hijmans, 2017) and additional climatic and biophysical variables from the ENVIREM database (Title & Bemmels, 2018). Each point (or bar) represents the strength of the relationship between genetic distance and a single environmental variable.

**Supplementary Fig. S7.**
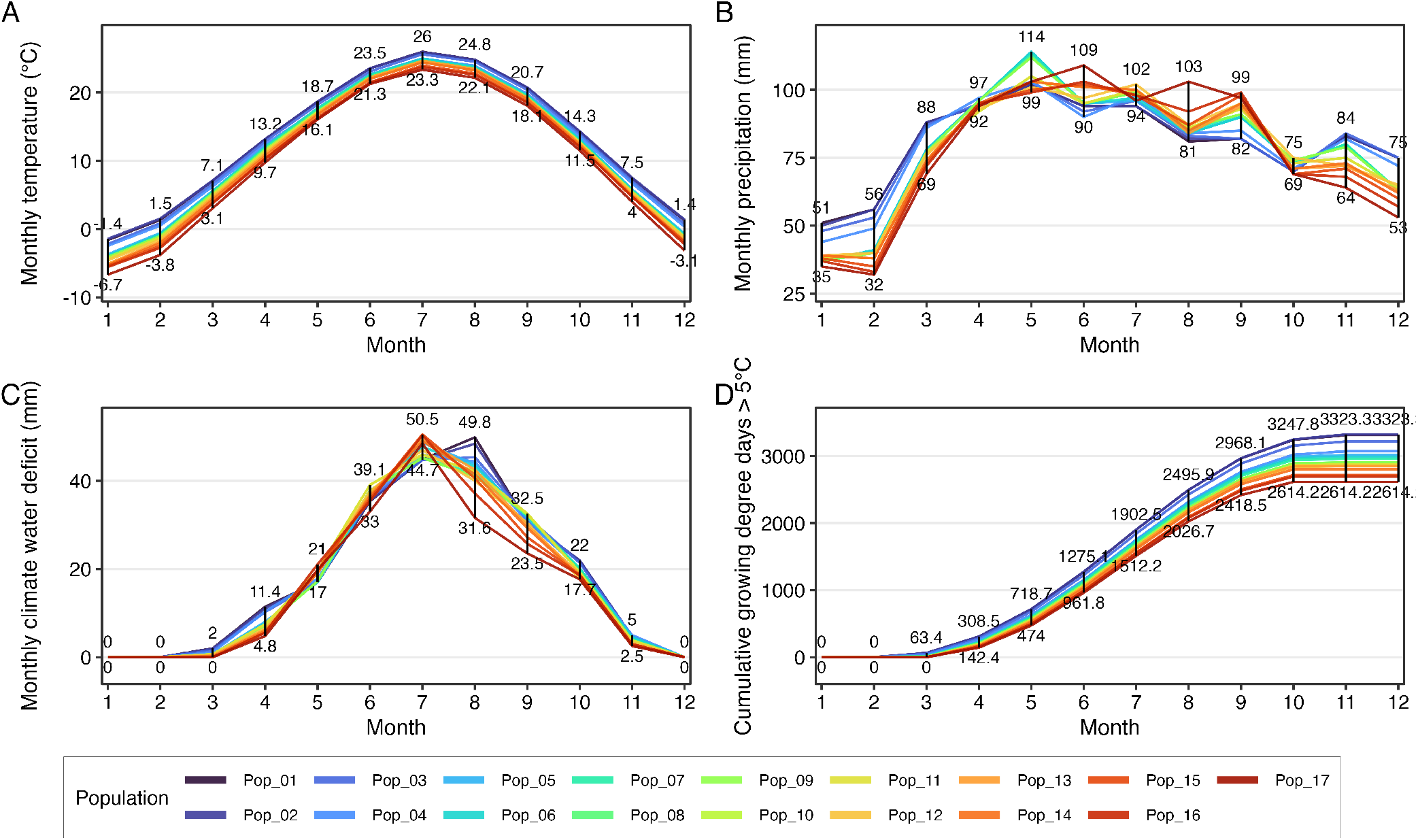
Monthly climatic profiles of study populations. Seasonal variation in four climatic variables across study populations: **(A)** Monthly temperature (°C). **(B)** Monthly precipitation (mm). **(C)** monthly climate water deficit (mm). **(D)** cumulative growing degree days above 5°C. Each colored line represents one population. For each month, black vertical bars indicate the observed range across populations, spanning the minimum and maximum values, and the numerical annotations indicate these extrema. Together, these panels illustrate both the shared seasonal climatic paiern and the extent of environmental heterogeneity among populations.

**Supplementary Fig. S8.**
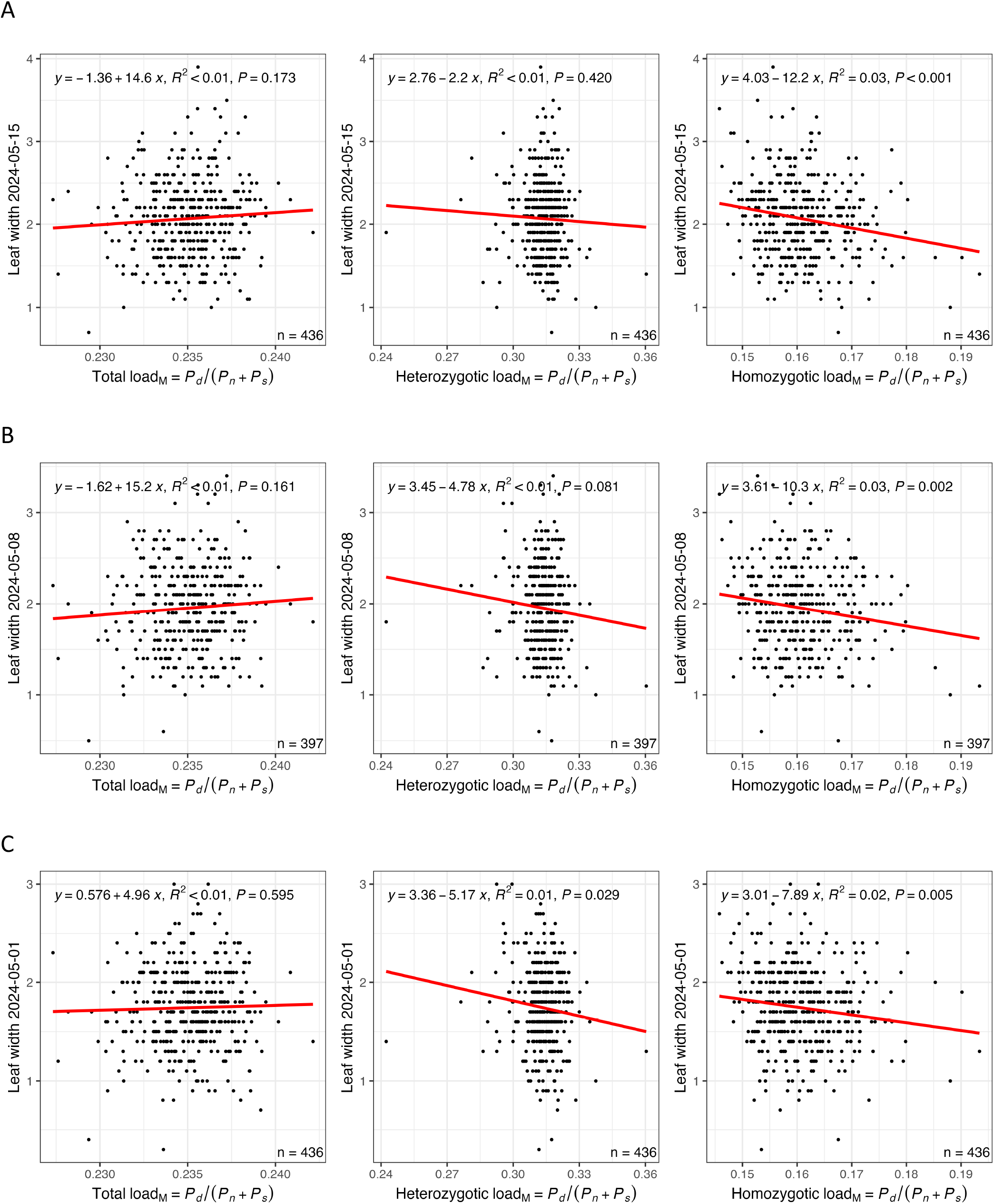
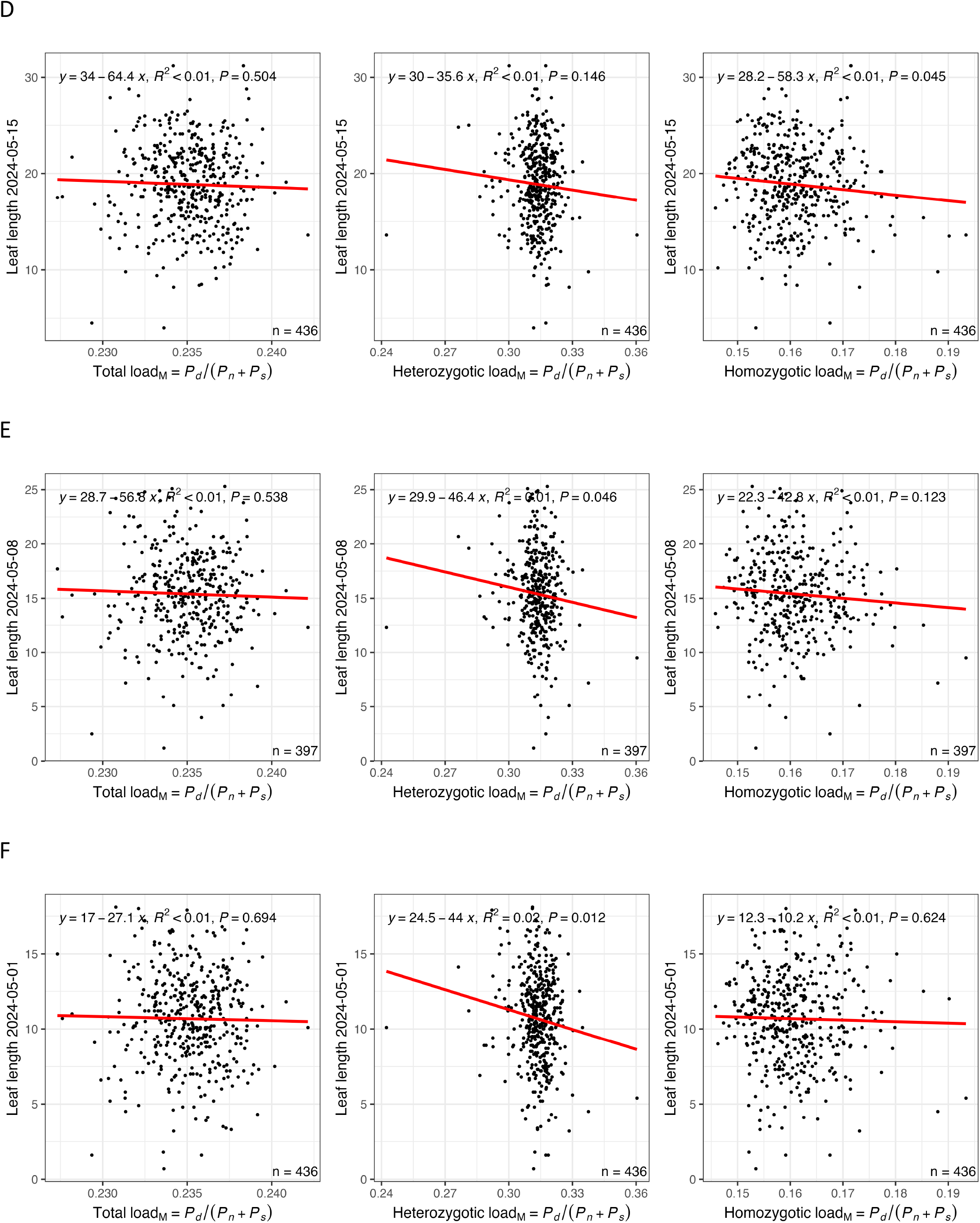
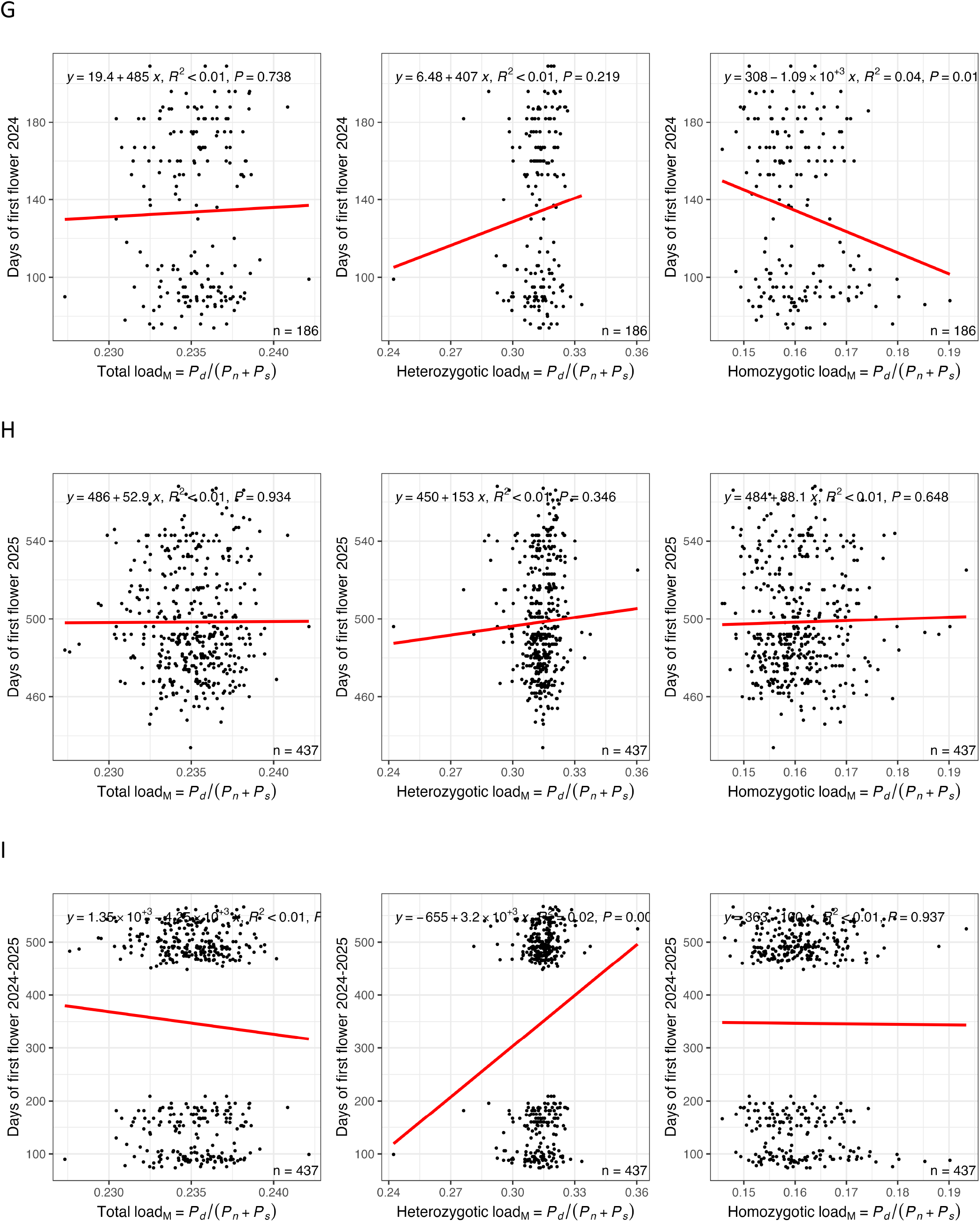
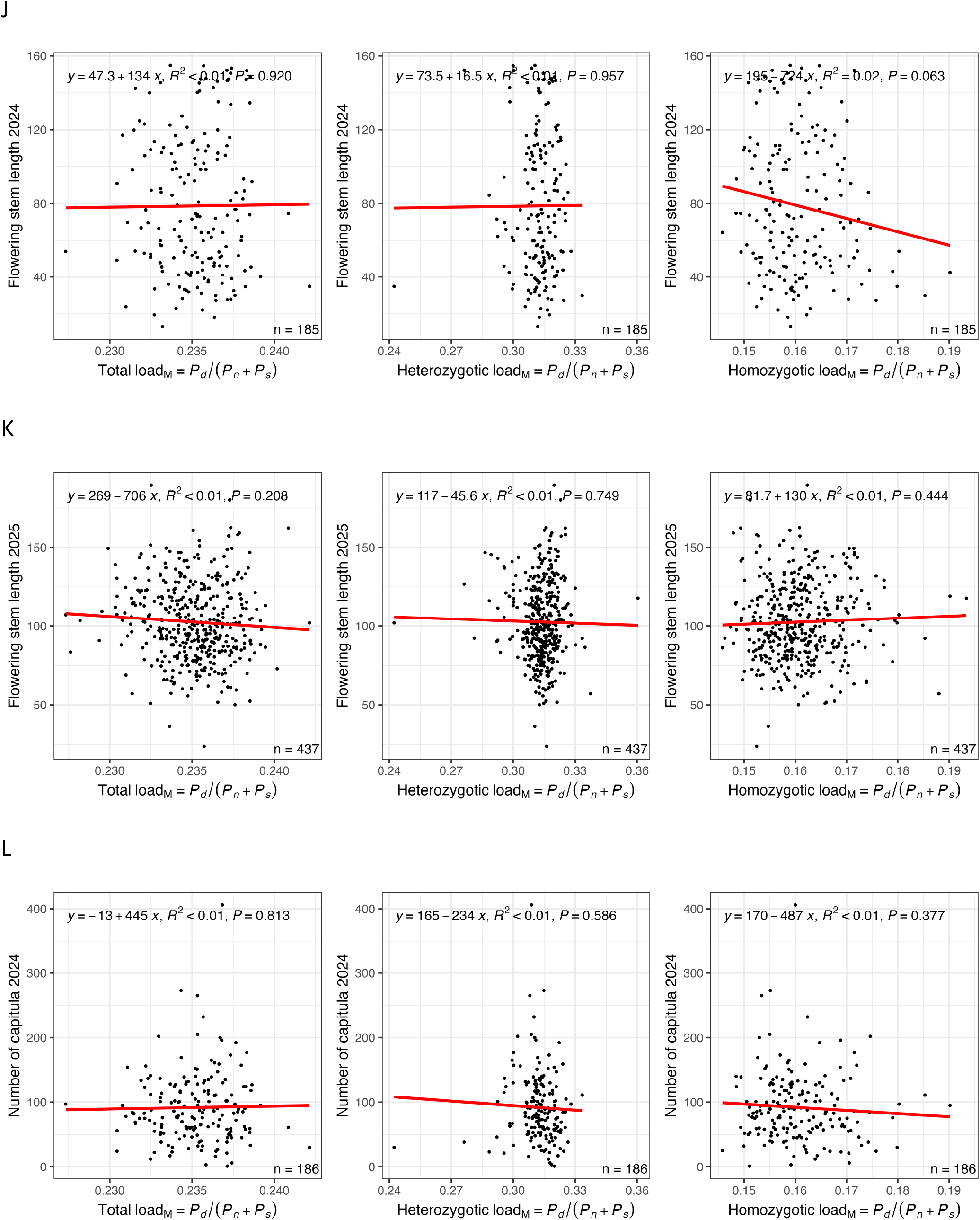
Individual genetic load and its association with phenotypic variation.

**Supplementary Fig. S9.**
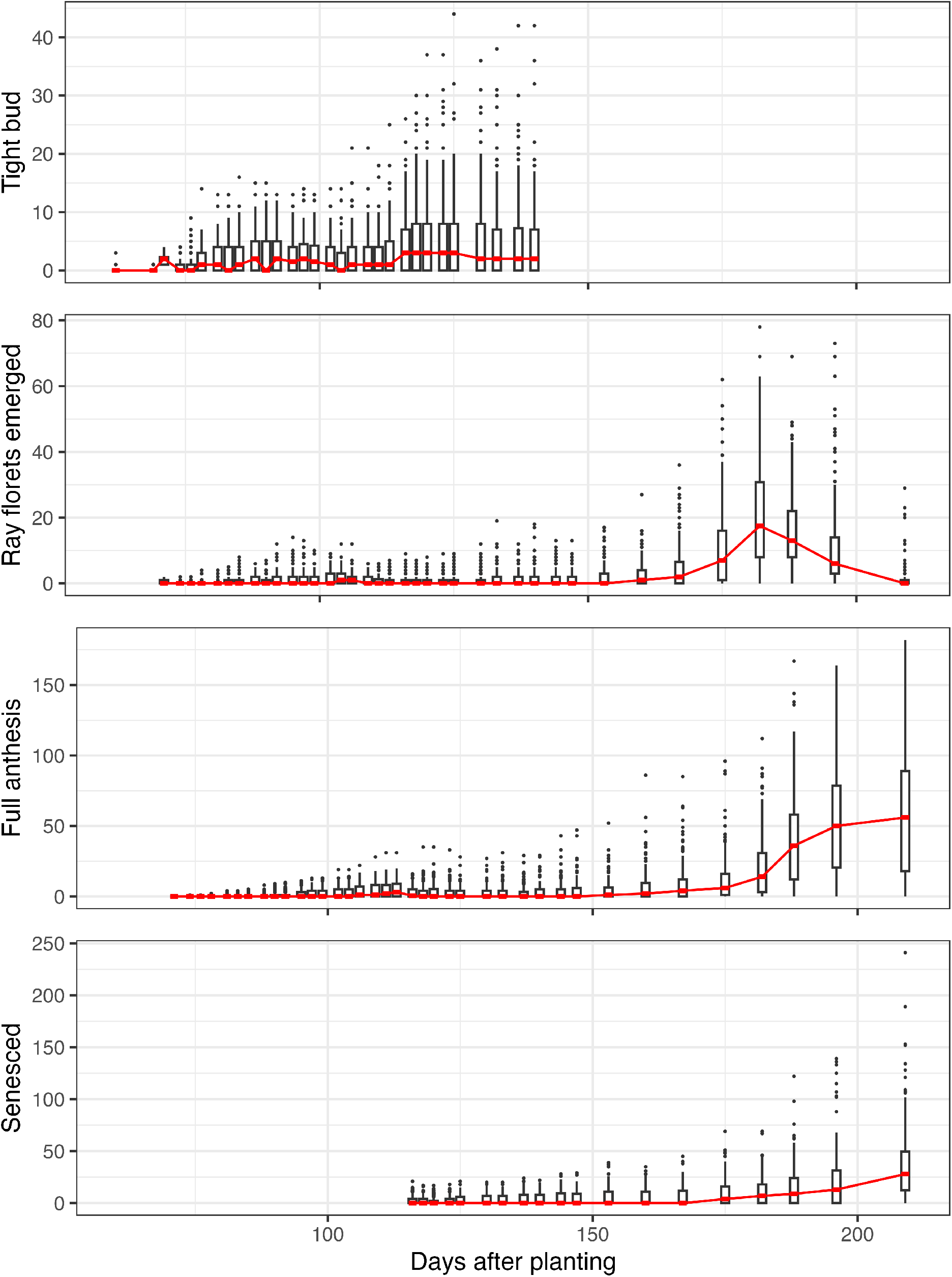
Documentation of reproductive traits in 2024.

**Supplementary Fig. S10.**
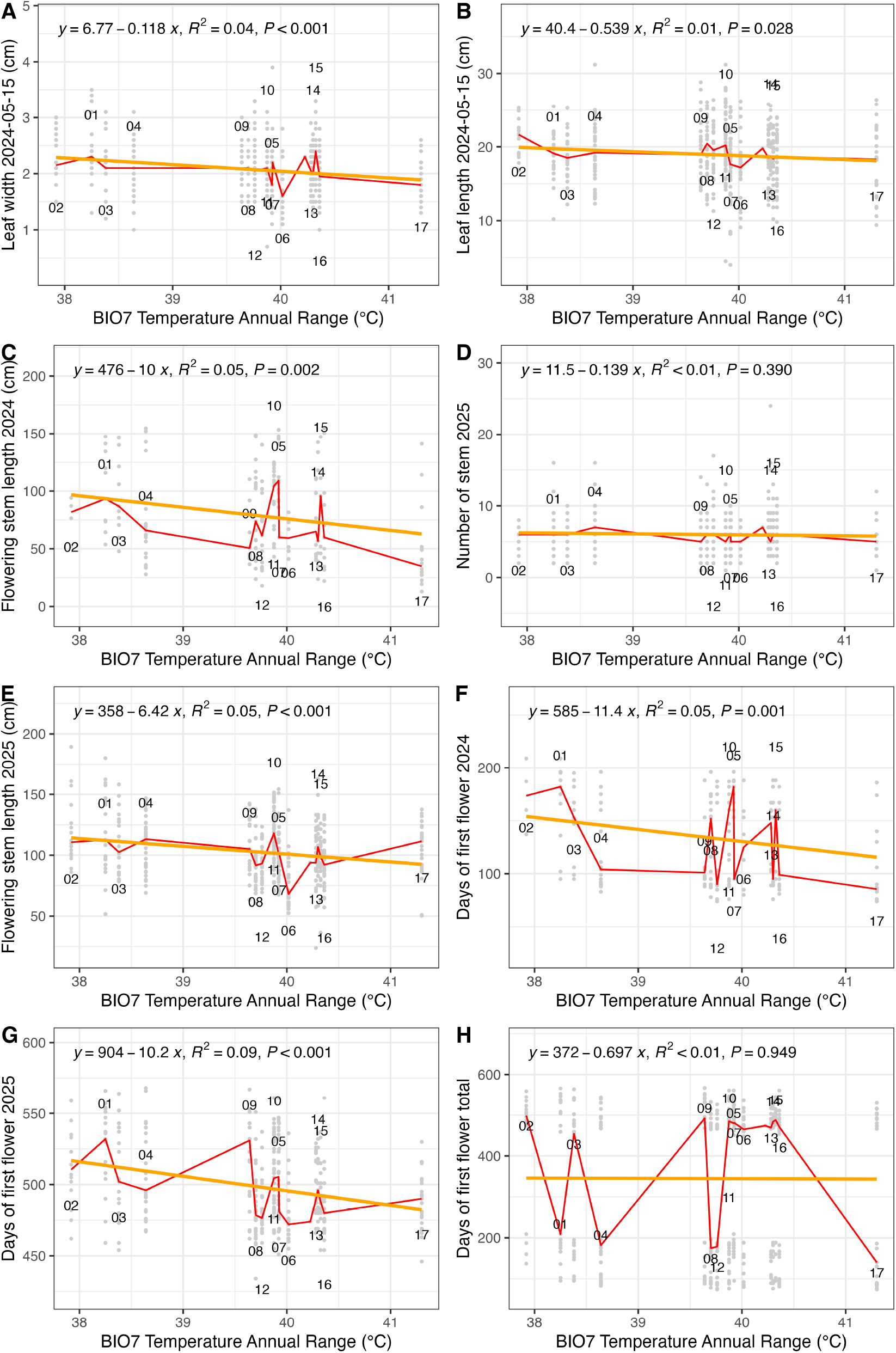
Phenotypic clines along BIO7 Temperature Annual Range. **(A)** Leaf width measured on 2024-05-15, which was the date shortly before the first tier of plants started to flower. The measured was taken from the longest leaf of a plant. **(B)** Leaf length measured on 2024-05-15. The measured was taken from the longest leaf of a plant. **(C)** Flowering stem length in 2024. The measured was the terminal length of the flower stem. **(D)** Number of stems in 2025. The measured was the taken on the date when the first disk flower in a capitulum opened. **(E)** Flowering stem length in 2025. The measured was the terminal length of the flower stem. **(F)** Days of the first flower opened in 2024. **(G)** Days of the first flower opened in 2025. **(H)** Absolute days until the first disk flower opened in a capitulum in the two years of the experiment.

