## Supplementary Information for "Genomic Consequences of Altered Flood Regimes: Gene Flow and Adaptation in a Threatened Floodplain Plant"

### Extended Materials and Methods

#### Reference genome sequencing and assembly

##### High-molecular-weight DNA extraction

Young, healthy leaf tissue was harvested from Boltonia_435 (Gilbert Lake 1997, Pop_04; [SAMN46503941](https://dataview.ncbi.nlm.nih.gov/object/55379665) = SAMN47833999), flash frozen in liquid nitrogen, and stored at −80 °C. High-molecular-weight genomic DNA was first isolated from 5 g of frozen leaf tissue using **PacBio plant nuclei isolation protocol (Workman et al., 2018)**, then a general CTAB DNA extraction protocol (Allen, Flores-Vergara, Krasynanski, Kumar, & Thompson, 2006). DNA integrity was confirmed by agarose gel electrophoresis and capillary electrophoresis, and DNA concentration and purity were assessed using spectrophotometry and fluorometry.

##### PacBio HiFi library preparation and Revio sequencing

SMRTbell libraries were constructed from HMW DNA using PacBio’s standard HiFi workflow for the **Revio** system. Libraries were size-selected on a **Sage Pippin HT** to enrich for long inserts, and library fragment distributions were checked on the Femto Pulse prior to sequencing. HiFi sequencing was carried out on a **PacBio Revio platform** at the **Genome Technology Access Center at the McDonnell Genome Institute at Washington University School of Medicine (GTAC@MGI; St. Louis, MO, USA)**.

##### Hi-C library preparation and sequencing

To obtain haplotig phasing and chromosome-scale scaffolding information for the Boltonia_435 genome, Hi-C libraries were prepared from isolated nuclei (see high-molecular-weight DNA extraction) using the Proximo Hi-C Plant kit (Phase Genomics, Seattle, WA, USA) according to the manufacturer’s instructions. The restriction enzyme cutting sites were GATC, CTNAG, TTAA, GANTC. The Hi-C libraries were sequenced using 2 × 150 bp reads on one lane of a 10B flow cell of Illumina NovaSeq X plus platform at Novogene Corporation Inc. (Sacramento, CA, USA). Sequencing yielded approximately 250 million read pairs, providing sufficient depth to generate a high-resolution chromatin contact map.

##### *De novo* assembly and scaffolding

*De novo* whole genome assembly was performed with hifiasm v0.20.0 in Hi-C mode (Cheng, Concepcion, Feng, Zhang, & Li, 2021). We ran hifiasm with an expected haploid genome size of 500 Mb (--hg-size 500m), enabling dual scaffolding with Hi-C reads. The final run produced two complete Hi-C–integrated assemblies: primary and alternative haplotigs.

Hi-C contact data were used to scaffold the assembly to the chromosome scale. Raw Hi-C read pairs were adapter-trimmed with Cutadapt (Martin, 2011), then aligned to the assembly and processed with Juicer 1.6 (-S early) (Durand, Shamim, et al., 2016). Scaffolding was performed with 3D-DNA 201008 (Dudchenko et al., 2017). The resulting scaffolds were manually curated in Juicebox Assembly Tools (Durand, Robinson, et al., 2016). Curated assemblies were exported from Juicebox. Finally, the two haplotigs were compared and any large-scale discrepancies were evaluated; only structural differences consistently supported by Hi-C signal were retained.

##### Contig Assembly curation and quality assessment

Prior to assembly, we used Jellyfish (Marçais & Kingsford, 2011) **and GenomeScope (Ranallo-Benavidez, Jaron, & Schatz, 2020)** to obtain initial estimates of genome size, sequencing depth, and heterozygosity directly from the PacBio HiFi read k-mer spectrum. After *de novo* assembly and Hi-C–based scaffolding, we performed multiple complementary assessments of assembly quality. First, we used **Merqury** to evaluate consensus accuracy and completeness based on k-mers derived from an independent Illumina short-read dataset. Illumina reads were quality-trimmed and used to generate a trusted k-mer set, which was then compared to the assembled genome to calculate the Merqury QV (phred-scaled consensus accuracy), k-mer completeness, and the fraction of k-mers present only in the reads or only in the assembly. We further evaluated contiguity and structural features of the assembly using **QUAST**.

To characterize the relationship between haplotypic representations produced by hifiasm, we aligned the primary and alternate assemblies using **mummer4** (nucmer). Whole-genome alignments between haplotigs were inspected to identify large-scale structural differences, regions of high divergence, and potential assembly inconsistencies. Finally, we assessed gene-space completeness using **BUSCO** with embryophyta_odb10, eudicots_odb10 database. Together, the GenomeScope, Merqury, QUAST, mummer4, and BUSCO results indicated that the Boltonia_435 assembly was of high quality and suitable to serve as the reference genome for downstream read mapping, variant calling, and population genomic analyses.

##### Gene annotation

Repetitive elements were identified and masked prior to gene prediction, with repeat annotation performed separately for each haplotype assembly. For each haplotype FASTA, we built a RepeatModeler database and generated a de novo repeat library with LTR discovery enabled, and then used the resulting haplotype-specific library to soft-mask repeats in the corresponding assembly with RepeatMasker. Structural gene annotation was then conducted with BRAKER3 using the repeat-masked genome and protein homology evidence from a curated Viridiplantae protein set.

#### Whole-genome resequencing

##### library preparation and sequencing

All fresh and herbarium leaf tissues were processed by the same CTAB-based DNA extraction protocol (Allen et al., 2006). Then, Illumina libraries were prepared using the **NEBNext Ultra II FS DNA Library Prep Kit** (New England Biolabs), following the manufacturer’s protocol. Fragmentation and size-selection settings were chosen to produce libraries with final (complete library construct) size distribution of approximately **400–600 bp**.

Adapters and indices were added using the **NEB Unique Dual Index (UDI) system**, generating uniquely dual-indexed libraries for each individual. Initial library quantification was performed with the **NEB Library Quantification Kit** on a **Bio-Rad CFX96** real-time PCR system, following the manufacturer’s instructions. Libraries were then pooled equimolarly based on qPCR-derived concentrations. The pooled library was **re-quantified** using the same NEB Library Quantification Kit and evaluated on an **Agilent TapeStation** using the **HS1000 kit** to verify fragment size distribution, absence of adapter dimers, and overall library integrity.

Sequencing was carried out at **Novogene Corporation Inc. (Sacramento, CA, USA)** on an Illumina NovaSeq X plus platform. We targeted at least **10× genome coverage per individual**. After each sequencing run, per-sample yield and coverage were assessed based on read counts and mapping statistics. **Samples that did not reach the target coverage (e.g., due to low library yield or under-representation in the pool) were resubmitted for additional sequencing** until their combined data met or exceeded the desired ~10× coverage threshold.

##### Variant site calling

We used the GATK best practice pipeline to generate multiple genome-wide, multi-sample VCF datasets tailored to specific analyses (Poplin et al., 2017). Raw paired-end Illumina whole-genome resequencing reads were first trimmed with Cutadapt (Martin, 2011) to remove adapter contamination using standard TruSeq adapter sequences. Trimmed reads were aligned to the Boltonia decurrens Boltonia_435 reference genome using BWA-MEM (Li & Durbin, 2009). PCR/optical duplicates were marked with GATK MarkDuplicates, after which we removed unmapped reads, secondary and supplementary alignments, and duplicates, and further restricted to high-confidence alignments by filtering for MAPQ ≥ 30; the resulting BAMs were re-indexed and standardized with complete read-group information using GATK AddOrReplaceReadGroups. Variants were then called per sample using GATK HaplotypeCaller in GVCF mode (-ERC GVCF) to generate one GVCF per sample, and joint genotyping was performed by aggregating GVCFs into interval-based GenomicsDB workspaces (GATK GenomicsDBImport) followed by multi-sample genotyping with GATK GenotypeGVCFs; interval VCFs were subsequently combined into a genome-wide multi-sample VCF.

##### VCF datasets

For assessment of species identity and hybridization, we compiled a dataset including all *Boltonia* species and the outgroup *Chloracantha spinosa* (prefix “Ball”), whereas analyses of population structure and functional genomics focused on *B. decurrens* only (prefix “Bdecurrens”). After joint genotyping, we applied a consistent baseline filtering workflow to each multi-sample VCF: SNPs and indels were hard-filtered separately using site-level thresholds (SNPs: QD < 2.0, FS > 60.0, MQ < 40.0, MQRankSum < −12.5, ReadPosRankSum < −8.0, SOR > 3.0; indels: QD < 2.0, FS > 200.0, ReadPosRankSum < −20.0, SOR > 10.0) and then concatenated, after which we retained only biallelic variants, masked low-confidence genotypes (DP < 3, DP > 50, or GQ < 10), recalculated INFO fields (AN, AC, AF, NS), removed sites with >10% missing genotypes across samples, and standardized ploidy to diploid before exporting the final VCF.

For the “Ball” dataset, because interspecific mapping rates can vary widely and inflate missingness in the “Ball” dataset, we applied an additional filtering step: we generate a subset VCF, including only non-*Boltonia decurrens* samples. Then, we only kept variant sites with ≤5% missing genotypes (F_MISSING ≤ 0.05), extracted the retained coordinates (CHROM and POS), and then applied this site list to the full “Ball” dataset. For phylogeny and admixture graph inference, a more stringent missing rate filter of 0.03 was applied to all remaining variant sites. Then, the dataset was LD pruned (PLINK --indep-pairwise 100kb 0.8) as “Ball_FMISSING003_LD” (625,566 variant sites). For analyses related to admixed graph, hybridization, introgression, we keep only non-missing sites in *B. apalachicolensis* (outgroup); then a more stringent missing rate filter of 0.01 was applied to all remaining variant sites. This dataset was called “Dsamples.B484_B485_nomiss. FMISS001” (170,217 variant sites). For speciation-positive selection scans, only pure *B. asteroides* and *B. decurrens* were kept; then a more stringent missing rate filter of 0.05 was applied to all remaining variant sites. This dataset was called “asteroides_samples” (1,538,608 variant sites).

For the “Bdecurrens” dataset, we additionally performed genotype imputation using Beagle v. 27Feb25.75f to generate “Bdecurrens_IMPUTATED” (9,957,183 variant sites), which was used for runs of homozygosity (ROH) and window-based summaries of nucleotide diversity (π) and Tajima’s D; we then filtered to minor allele frequency (MAF) >0.01 to create “Bdecurrens_IMPUTATED_MAF” (6,950,268 variant sites) for GWAS, genetic differentiation, and landscape genomic analyses. Finally, we generated analysis-specific LD-pruned derivatives, including a mildly pruned dataset (PLINK --indep-pairwise 100kb 0.8) for PCA, ADMIXTURE, and F_ST analyses (“Bdecurrens_IMPUTATED_MAF_mildLD”; 1,863,350 variant sites) and a higher-LD-retained dataset used for fastsimcoal2 demographic modeling (“Bdecurrens_IMPUTATED_highLD”).

### Supplementary Tables

**Supplementary Table S1.** *Boltonia decurrens* genome assembly summary

|  | Primary* | Alternative* |
| --- | --- | --- |
| HiFi Read coverage | Homozygous regions: 88, Heterozygous regions: 43 | |
| Number of contigs | 172 | 60 |
| Contig N50 (bp) | 47,358,366 | 46,711,460 |
| Longest contigs | 74,295,378 | 80,471,637 |
| Number of scaffolds | 169 | 56 |
| Scaffold N50 (bp) | 47,358,366 | 46,926,256 |
| Size of final assembly (bp) | 448,103,187 | 449,760,986 |
| Base pair QV | 38.6164 | 38.7498 |
| Base pair QV (both) | 38.6827 | |
| k-mer completeness | 81.5472% | 81.8373% |
| k-mer completeness (both) | 99.3233% | |
| BUSCO  eudicots_odb10 (2326 genes) | C: 95.6% [S: 91.9%, D: 3.7%],  F: 0.6%, M: 3.8% | C: 95.4% [S: 92.0%, D: 3.4%],  F: 0.8%, M: 3.8% |
| BUSCO  embryophyta_odb10 (1614 genes) | C: 97.7% [S: 94.5%, D: 3.2%],  F: 0.6%, M: 1.7% | C: 97.5% [S: 94.4%, D: 3.1%],  F: 0.6%, M: 1.9% |
| Assigned chromosomes | 97.692% | 98.884% |
| # N's | 128,830 | 752,087 |
| # N's per 100 kbp | 28.75 | 167.22 |
| GC level | 34.46% | 34.46% |
| Repeated elements | 57.54% | 57.09% |
| Retroelements | 15.33% | 16.73% |
| LTR elements (Ty1/Copia) | 5.67% | 6.10% |
| LTR elements (Gypsy/DIRS1) | 6.84% | 7.38% |
| DNA transposons | 2.39% | 2.56% |

*Representing the two haploid genomes of a diploid organism

**Supplementary Table S2.** Candidate genes for adaptive evolution in floodplain habitats

| Gene_ID | Gene_Symbol | Protein_Name | Uniprot_Accession | Potential_Function |
| --- | --- | --- | --- | --- |
| g1629 | LPEAT2 | Lysophospholipid acyltransferase LPEAT2 | Q8S8S2 | Enzyme controlling phospholipid acyl composition and membrane lipid homeostasis. |
| g2481 | Xyl2 | Beta-xylosidase/alpha-L-arabinofuranosidase 2 | A5JTQ3 | Cell-wall hydrolase modifying hemicellulose and arabinoxylan side chains. |
| g2850 | COP1 | E3 ubiquitin-protein ligase COP1 | P43254 | Master E3 ligase regulator of light signaling and photomorphogenesis. |
| g3862 | THO6 | THO complex subunit 6 | Q8L4M1 | mRNA-processing and export factor that also helps repress ABA signaling via an E3 ligase complex. |
| g5249 | PRE6 | Transcription factor PRE6 | Q8GW32 | bHLH transcriptional regulator modulating shade-avoidance and light responses. |
| g12067 | RCD1 | Inactive poly [ADP-ribose] polymerase RCD1 | Q8RY59 | Regulator of oxidative and hormonal stress responses integrating multiple abiotic stress signals. |
| g12106 | SIS8 | Probable serine/threonine-protein kinase SIS8 | Q9C9U5 | Kinase linking salt tolerance with sugar signaling during early seedling development. |
| g12166 | ASP5 | Aspartate aminotransferase, chloroplastic | P46248 | Chloroplastic aminotransferase central to nitrogen and carbon metabolism. |
| g20122 | PCMP-H40 | Pentatricopeptide repeat-containing protein At1g11290, chloroplastic | Q3E6Q1 | Chloroplast PPR protein controlling RNA editing of NDH and rpoB, affecting NDH activity. |
| g22695 | CAD1 | Probable cinnamyl alcohol dehydrogenase 1 | P42495 | Lignin-pathway enzyme producing monolignol alcohols for cell wall strengthening. |
| g25346 | PCMP-H59 | Pentatricopeptide repeat-containing protein At5g50990 | Q9FI49 | Chloroplast PPR protein likely involved in organellar RNA processing and electron transport. |
| g25347 | NRP | DCD domain-containing protein NRP | Q8RXN8 | Modulator that enhances ABA responses during abiotic and biotic stress. |
| g26906 | FTIP3 | FT-interacting protein 3 | Q9M2R0 | Regulator of shoot apical meristem stem-cell balance and shoot architecture. |
| g30468 | At5g60760 | P-loop NTPase domain-containing protein LPA1 homolog 1 | Q9FJH9 | Regulator of seed phytic acid levels and phosphorus storage. |
| g30917 | MPV17 | Mitochondrial inner membrane protein Mpv17 | Q2KIN6 | Mitochondrial channel controlling membrane potential, dNTP pools, and ROS homeostasis. |
| g30919 | ITN1 | Ankyrin repeat-containing protein ITN1 | Q9C7A2 | ABA- and ROS-related regulator contributing to salt and osmotic stress tolerance. |
| g31612 | PLDZETA1 | Phospholipase D zeta 1 | Q9LRZ5 | Phospholipase involved in root-hair development and phosphorus-starvation responses. |

**Supplementary Table S3.** Candidate genes for flowering time variation

| Gene_ID | Gene_Symbol | Protein_Name | Uniprot_Accession | Potential_Function |
| --- | --- | --- | --- | --- |
| g1365 | SKP1B | SKP1-like protein 1B | Q9FHW7 | Involved in ubiquitination and subsequent proteasomal degradation of target proteins. Together with CUL1, RBX1 and a F-box protein, it forms a SCF E3 ubiquitin ligase complex. The functional specificity of this complex depends on the type of F-box protein. In the SCF complex, it serves as an adapter that links the F-box protein to CUL1. SCF(UFO) is required for vegetative and floral organ development as well as for male gametogenesis. SCF(TIR1) is involved in auxin signaling pathway. SCF(COI1) regulates responses to jasmonates. SCF(EID1) and SCF(AFR) are implicated in phytochrome A light signaling. SCF(ADO1), SCF(ADO2), SCF(ADO3) are related to the circadian clock. SCF(ORE9) seems to be involved in senescence. SCF(EBF1/EBF2) may regulate ethylene signaling. Plays a role during embryogenesis and early postembryonic development, especially during cell elongation and division. Contributes to the correct chromosome segregation during tetrad formation. {ECO:0000269\|PubMed:10398681, ECO:0000269\|PubMed:12970487, ECO:0000269\|PubMed:14688296}. |
| g25292 | CIPK5 | CBL-interacting serine/threonine-protein kinase 5 | Q9LEU7 | CIPK serine-threonine protein kinases interact with CBL proteins. Binding of a CBL protein to the regulatory NAF domain of CIPK protein lead to the activation of the kinase in a calcium-dependent manner (By similarity). {ECO:0000250}. |
| g28510 | ETO1 | Ethylene-overproduction protein 1 | O65020 | Essential regulator of the ethylene pathway, which acts by regulating the stability of 1-aminocyclopropane-1-carboxylate synthase (ACS) enzymes. May act as a substrate-specific adapter that connects ACS enzymes, such as ACS5, to ubiquitin ligase complexes, leading to proteasomal degradation of ACS enzymes. {ECO:0000269\|PubMed:15118728, ECO:0000269\|PubMed:16091151, ECO:0000269\|PubMed:18808454}. |
| g28526 | FLX | Protein FLC EXPRESSOR | F4IMQ0 | Transactivator required for FRIGIDA-mediated activation of FLC and for the expression of the repressors of flowering AGL27/MAF1 and AGL31/MAF2. Does not alter expression of known activators of FLC expression. Not required for the photoperiod response or for the vernalization response. Not redundant with FLXL4. {ECO:0000269\|PubMed:18156133, ECO:0000269\|PubMed:21282526, ECO:0000269\|PubMed:23864009}. |

**Supplementary Table S4.** Candidate genes for stem length variation

| Gene_ID | Gene_Symbol | Protein_Name | Uniprot_Accession | Potential_Function |
| --- | --- | --- | --- | --- |
| g1954 | At1g07650 | Probable LRR receptor-like serine/threonine-protein kinase At1g07650 | C0LGE0 | LRR receptor-like kinase; plausible regulator of hormone or peptide signaling influencing cell expansion and meristem activity. |
| g16073 | Os07g0190000 | Probable 1-deoxy-D-xylulose-5-phosphate synthase 2, chloroplastic | Q6YU51 | Catalyzes the acyloin condensation reaction between C atoms 2 and 3 of pyruvate and glyceraldehyde 3-phosphate to yield 1-deoxy-D-xylulose-5-phosphate (DXP). Is a limiting enzyme for plastidic isoprenoid biosynthesis and essential for chloroplast development (By similarity). {ECO:0000250}. |
| g20276 | ERF003 | Ethylene-responsive transcription factor ERF003 | Q94AW5 | Probably acts as a transcriptional activator. Binds to the GCC-box pathogenesis-related promoter element. May be involved in the regulation of gene expression by stress factors and by components of stress signal transduction pathways (By similarity). {ECO:0000250}. |
| g20277 | AHA9 | ATPase 9, plasma membrane-type | Q42556 | The plasma membrane H(+) ATPase of plants and fungi generates a proton gradient that drives the active transport of nutrients by H(+)-symport. The resulting external acidification and/or internal alkinization may mediate growth responses. |
| g20278 | DCL4 | Endoribonuclease Dicer homolog 4 | A7LFZ6 | Involved in the RNA silencing pathway. Cleaves double-stranded RNA to produce small interfering RNAs (siRNAs) which target the selective destruction of complementary RNAs. Required for the production of 21 nucleotide siRNAs. Regulates shoot apical meristem (SAM) initiation and maintenance, leaf polarization and lemma polarity through the trans-acting siRNAS (ta-siRNAs) pathway, which probably modulate the expression of the ARF2, ARF3, ARF4, ARF14 and ARF15 genes. Can process endogenous 21 nucleotide siRNAs derived from an imperfect inverted repeat. May not be involved in microRNAs (miRNAs) production. {ECO:0000269\|PubMed:17804793, ECO:0000269\|PubMed:17905898}. |
| g30592 | KIN7K | Kinesin-like protein KIN-7K, chloroplastic | F4J8L3 | Kinesin-like protein; cytoskeleton/organelle dynamics may impact cell elongation and organ growth, though mechanism is less specific than AHA9/DXS2. |

**Supplementary Table S5.** Candidate genes for stem number variation

| Gene_ID | Gene_Symbol | Protein_Name | Uniprot_Accession | Potential_Function |
| --- | --- | --- | --- | --- |
| g7188 | LTL1 | GDSL esterase/lipase LTL1 | Q9M8Y5 | Involved in the mechanisms of salt tolerance (PubMed:16930315). Mediates resistance to LiCl and NaCl (PubMed:16930315). {ECO:0000269\|PubMed:16930315}. |
| g23772 | RIN13 | RPM1 interacting protein 13 | Q9SK71 | Resistance protein interactor which positively enhances RPM1-mediated resistance to necrotrophic bacterial pathogens Pseudomonas syringae pv. tomato DC3000 harboring type III effector protein AvrRpm1 or AvrB, but prevents the hypersensitive response (HR) controlled by RPM1 (PubMed:15722472). Together with ARF1, promotes leaf senescence and cell death, probably by facilitating the translocation of ARF1 into the nucleus, and activates ROS-related enzymes (e.g. POD, CAT and SOD) (PubMed:32446355). {ECO:0000269\|PubMed:15722472, ECO:0000269\|PubMed:32446355}. |
| g23775 | IAA13 | Auxin-responsive protein IAA13 | Q38831 | Aux/IAA proteins are short-lived transcriptional factors that function as repressors of early auxin response genes at low auxin concentrations. Repression is thought to result from the interaction with auxin response factors (ARFs), proteins that bind to the auxin-responsive promoter element (AuxRE). Formation of heterodimers with ARF proteins may alter their ability to modulate early auxin response genes expression. {ECO:0000269\|PubMed:12036262}. |
| g33823 | RAB7 | Ras-related protein Rab7 | Q43463 | Protein transport. Probably involved in vesicular traffic (By similarity). {ECO:0000250}. |
| g35152 | CHS3 | Chalcone synthase 3 | P48392 | The primary product of this enzyme is 4,2',4',6'-tetrahydroxychalcone (also termed naringenin-chalcone or chalcone) which can under specific conditions spontaneously isomerize into naringenin. |
